# Rapid evolution at the Drosophila telomere: transposable element dynamics at an intrinsically unstable locus

**DOI:** 10.1101/782904

**Authors:** Michael P McGurk, Anne-Marie Dion-Côté, Daniel A Barbash

**Author notes:** CORRESPONDING AUTHOR Daniel A. Barbash, Department of Molecular Biology and Genetics, 403 Biotechnology Building, Cornell University, Ithaca, NY 14853. These authors contributed equally to this work (Michael P McGurk, Anne-Marie Dion-Côté).

## Abstract

Drosophila telomeres have been maintained by three families of active transposable elements (TEs), *HeT-A, TAHRE* and *TART*, collectively referred to as HTTs, for tens of millions of years, which contrasts with an unusually high degree of HTT interspecific variation. While the impacts of conflict and domestication are often invoked to explain HTT variation, the telomeres are unstable structures such that neutral mutational processes and evolutionary tradeoffs may also drive HTT evolution. We leveraged population genomic data to analyze nearly 10,000 HTT insertions in 85 *D. melanogaster* genomes and compared their variation to other more typical TE families. We observe that occasional large-scale copy number expansions of both HTTs and other TE families occur, highlighting that the HTTs are, like their feral cousins, typically repressed but primed to take over given the opportunity. However, large expansions of HTTs are not caused by the runaway activity of any particular HTT subfamilies or even associated with telomere-specific TE activity, as might be expected if HTTs are in strong genetic conflict with their hosts. Rather than conflict, we suggest instead that distinctive aspects of HTT copy number variation and sequence diversity largely reflect telomere instability, with HTT insertions being lost at much higher rates than other TEs elsewhere in the genome. We extend previous observations that telomere deletions occur at a high rate, and surprisingly discover that more than a third do not appear to have been healed with an HTT insertion. We also report that some HTT families may be preferentially activated by the erosion of whole telomeres, implying the existence of HTT-specific host control mechanisms. We further suggest that the persistent telomere localization of HTTs may reflect a highly successful evolutionary strategy that trades away a stable insertion site in order to have reduced impact on the host genome. We propose that HTT evolution is driven by multiple processes with niche specialization and telomere instability being previously underappreciated and likely predominant.

## INTRODUCTION

Transposable elements (TEs) are genomic parasites that can increase their copy number within genomes by a variety of transposition mechanisms. While this provides a replicative advantage to the TE, their mobilization has consequences to the host genome that include DNA double-strand breaks, disruption of open reading frames and regulatory elements, and the perturbation of gene expression (Bourque *et al*. 2018). The presence of dispersed repeats further provides substrates for ectopic recombination, permitting large scale and potentially lethal genome arrangements (Deininger *et al*. 2003). The resulting conflict between TEs and the host genome may progress in several ways.

The simplest models suggest that TE copy number may stabilize at an equilibrium between the fitness advantage that transposition confers to the TEs and the fitness costs imposed on the host genome (Charlesworth and Charlesworth 1983; Charlesworth and Langley 1989). This stabilization is now known to involve complex interactions with the host-defense piRNA system that utilizes small RNAs to repress TE activity (Lee and Langley 2010; Blumenstiel 2011; Kelleher *et al*. 2020). TEs may further alter these dynamics by adopting strategies that mitigate the mutational burden they impose on the genome without sacrificing their replicative success (Cosby *et al*. 2019). Some TEs exhibit insertion site preferences that restrict the set of loci into which they transpose and could in principle limit the potential for deleterious insertions (Sultana *et al*. 2017). Arthropod R-elements are an extreme example, as they only insert in the highly repeated ribosomal DNA (rDNA) genes (Eickbush 2002). Other models that incorporate inactivating mutations suggest saltatory dynamics, where TE families are successively replaced by more active subfamilies until the family is eventually lost from the genome (Le Rouzic *et al*. 2007). TE exaptation or domestication, whereby TE regulatory or coding sequences are co-opted by their host, is another path by which such conflicts may be resolved (Jangam *et al*. 2017). These instances do not typically impact the dynamics of the entire TE family, but rather preserve only a portion of a single TE insertion while the rest of the family independently lives or dies. As such, most examples of domesticated TEs have lost their transposition activity.

The telomeric TEs of Drosophila are a remarkable exception to how this conflict typically proceeds, where the transposition activity of several TE families performs a function essential for genome integrity. In most eukaryotes, telomeric DNA is comprised of simple repeats synthesized by telomerase that are assembled into a complex nucleoprotein structure. Telomeres serve two major roles: they protect chromosome ends from genetic attrition due to the end-replication problem, and they prevent chromosome ends from being recognized as DNA double-strand breaks, which can lead to chromosome fusions (de Lange 2009). In Diptera, the telomerase gene was lost (Mason *et al*. 2016) and in Drosophila its role in telomere elongation replaced by three non-LTR retrotransposons from the *jockey* clade: *HeT-A, TART* and *TAHRE* (herein collectively referred to as HTTs) (Levis *et al*. 1993; Abad *et al*. 2004). HTTs have several unique features compared to other non-LTR retrotransposons that reflect their specialized function (Mason *et al*. 2008; Pardue and DeBaryshe 2011; Arkhipova 2012). First, they are thought to exclusively insert at the ends of chromosomes, resulting in head-to-tail tandem arrays (Saint-Léandre *et al*. 2019; Pardue and DeBaryshe 2008). Second, they rely upon each other at several steps in their life cycles. *HeT-A* and *TARHE* in *D. melanogaster* carry promoters in their 3’ UTR that drive expression of their neighboring element, an innovation only possible because of HTT tandem arrangement (Danilevskaya *et al*. 1997; Maxwell *et al*. 2006). Additionally, *HeT-A* may provide telomere specificity to *TART* and *TAHRE* through its Gag-like protein encoded by ORF1 (Fuller *et al*. 2010). Finally, *HeT-A* is non-autonomous (Biessmann *et al*. 1992a; b; 1994) — it likely depends upon the reverse transcriptase encoded by *TAHRE* and/or *TART* for integration.

Despite being ancestral to Drosophila and having comprised the telomeres for ∼60 million years, prior work has uncovered extensive interspecific variation among HTTs, with new families and subfamilies having evolved across the genus and even among closely related species of the *melanogaster* subgroup (Villasante *et al*. 2008; Saint-Léandre *et al*. 2019). In addition, features typical of a family in one species, for example 3’-promoters or terminal repeats, may be absent in other species (Pardue and DeBaryshe 2011). This is a striking degree of variation given that HTTs are required for the fundamental role of protecting chromosome ends. These dramatic changes over the macroevolutionary scale must reflect a complex array of processes ongoing at the population level in ways distinct from more typical TEs. Yet, a comprehensive population genomic study of HTT variation remains to be undertaken, which could provide the power to delineate the forces shaping their contemporary evolution. We consider four potential forces shaping HTT variation at the population scale.

First, selection acting at the level of the organism has likely played an important role in both the evolution and dynamics of the HTTs, in ways distinct from more typical TEs. The HTTs comprise the Drosophila telomeres, which is often considered a paradigm of TE domestication, and should favor their long-term persistence and transposition activity. Organismal selection may also favor increased levels of domestication, bringing telomere elongation more tightly under host regulation. In addition, selection might promote new HTT variants that increase organismal fitness in other ways, for example by reducing telomere instability or by having pleiotropic beneficial effects on chromosome structure. Further, unlike other TEs, the genome is under pressure to regulate rather than wholly suppress their activity. Coordinating telomere elongation to prevent either complete erosion or overextension is a feature of more typical telomerase-based systems (Stewart *et al*. 2012; Zhao *et al*. 2014), and mutational studies have shown that Drosophila has recruited the piRNA and heterochromatin maintenance pathways typically involved in TE suppression into this role (Shpiz and Kalmykova 2012). Host factors that interact with the telomeres, such as the capping proteins that assemble onto them and the subtelomeric sequence which directly abut them, may have also evolved properties that regulate HTT activity (Raffa *et al*. 2011).

Second, selection can act on the HTTs themselves and place them in conflict with each other and with their host, because they are active transposons and thus replicative entities in their own right. Both the continual erosion of the telomeres and the potential fitness impacts of excessively long telomeres imply that space in the telomere is a limiting resource for which all HTT families are competing. HTT variants that are better able to increase in copy number will outcompete those that cannot, for example by escaping regulation by the host genome. Conflict between the HTTs and the host genome (or among the HTTs) is thus an ever present possibility. Indeed, like their feral cousins, studies using mutations in HTT regulators suggest that they are capable of taking over given the opportunity (Shpiz and Kalmykova 2012). Similar to the rapid evolution of the Drosophila HTTs (Villasante *et al*. 2007, 2008), some telomere-associated proteins show high evolutionary rates and signatures of positive selection (Raffa *et al*. 2011; Lee *et al*. 2017). These evolutionary analyses led Lee et al. to propose that HTTs are in genetic conflict with their Drosophila hosts (Lee *et al*. 2017). Complete resolution of conflict with the genome would likely require somehow separating the reverse transcription machinery that extends the telomere from the HTTs (Arkhipova 2012; Jangam *et al*. 2017; Markova *et al*. 2020).

Third, the dynamic and unstable nature of telomeres has the potential to shape HTT evolution. Telomeres continually experience terminal erosion due to the end-replication problem. In addition, their tandemly arrayed structure facilitates the amplification and deletion of sequence through unequal exchange events, and chromosome breaks near the terminus can cause complete loss of a telomere (Begun and Aquadro 1995; Langley *et al*. 2000; Kern and Begun 2008). The expansions, contractions, and deletions of telomeric sequence also have the potential to heighten the rate at which polymorphisms fix within HTT families and to increase the potential for the extinction of lineages, as observed recently in *D. biarmipes* (Saint-Léandre *et al*. 2019). The net effect of these processes is to facilitate rapid sequence change at telomeres in a manner driven by mutational processes rather than selection.

Fourth, the trade-off between reducing their impact on the genome versus inhabiting an unstable locus has likely also shaped HTT evolution (Markova et al. 2020). Compared to much of the genome, the telomere is a “safe harbor” that allows the HTTs to accumulate without disrupting essential host genes and also minimizes the potential effects of ectopic recombination (Cosby *et al*. 2019). The tension between this and the inherent instability of the telomeres disposing HTTs to high rates of deletion suggests that the persistence of this strategy reflects the outcome of an evolutionary trade-off. Indeed, we suggest that some of the peculiar features of HTTs, such as 3’-end promoters and head-to-tail arrangement, are likely adaptations to cope with this instability (Pardue and DeBaryshe 2008). Another possible outcome of evolutionary trade-offs (as well as of genetic conflict with other HTTs) would be to escape the telomere. Surprisingly, despite transposing to chromosome ends, *TART* and *TAHRE* each retain a potentially functional endonuclease domain (Casacuberta 2017), which raises the possibility that the HTTs could generate double-strand breaks and insert randomly as do other non-LTR retrotransposons. While limited surveys suggest that HTTs exclusively insert at telomeres in *D. melanogaster*, the *melanogaster* subgroup species *D. rhopaloa* has many copies outside the telomere, thus supporting the notion that HTTs can evolve the ability to escape from telomeres (Saint-Léandre *et al*. 2019).

The extent to which the HTTs vary among species should reflect the compounded effects and interaction among these evolutionary processes playing out over millions of years. However, the timescales involved and the complexity of this array of processes makes it challenging to understand which processes are responsible for which particular aspects of HTT rapid evolution. A clear picture of within-species HTT variation offers the opportunity to understand how these processes play out over shorter timescales where their impacts on HTT evolution can be more clearly distinguished. This requires examining telomere variation along with other active TEs in a large population sample. To date, two studies have presented telomere assembly solely in the reference genome of *D. melanogaster* (Saint-Léandre *et al*. 2019; George *et al*. 2006), while length variation has been assayed only in a few lines, or in a single population for only for a single site in *Het-A* (Siriaco et al. 2002; George et al. 2006; Wei et al. 2017).

Here, we leverage available population genomic data and the ConTExt pipeline (McGurk and Barbash 2018) to comprehensively analyze HTT sequence and copy number polymorphism in the Drosophila Global Diversity Lines (GDL) (Grenier *et al*. 2015). This dataset encompasses nearly 10,000 HTT insertions in 85 strains of *Drosophila melanogaster*, making it the largest study of telomere variation in Drosophila to date. We compare the HTTs to other TE families, including those that are similarly restricted to unstable genomic loci. Within this framework we ask whether the patterns of HTT copy number, organization, and sequence variation require explanations beyond the standard conflict between HTTs and the host genome and, if so, whether this reflects their remarkable symbiosis with the genome or is instead the natural consequence of occupying the unstable telomeres.

## RESULTS

### Analyzing telomere variation in the Global Diversity Lines using ConTExt

To explore HTT population variation, we leveraged the paired-end NGS data from the Global Diversity Lines, 85 stocks of *D. melanogaster* collected from five populations: Beijing, Ithaca, Netherlands, Tasmania, and Zimbabwe (Grenier *et al*. 2015). We employed the ConTExt pipeline (McGurk and Barbash 2018) to summarize telomere structure, which is composed of head-to-tail HTT tandem arrays (Figure 1A). In brief, ConTExt aligns repeat-derived reads to repeat-consensus sequences and uses paired-end information to identify the junctions between repeats and neighboring sequence (Figure 1B). In the case of HTT insertions, the neighboring sequence should generally be another HTT insertion or subtelomeric sequence (Figure 1A), and all such junctions should involve the intact 3’-end of a HTT. The range of 5’-truncations resulting from incomplete reverse transcription, which is frequent for all non-LTR retrotransposons, provides additional power to distinguish independent insertions, allowing us to identify insertions and estimate TE copy number within the telomeres.

**Figure 1.**
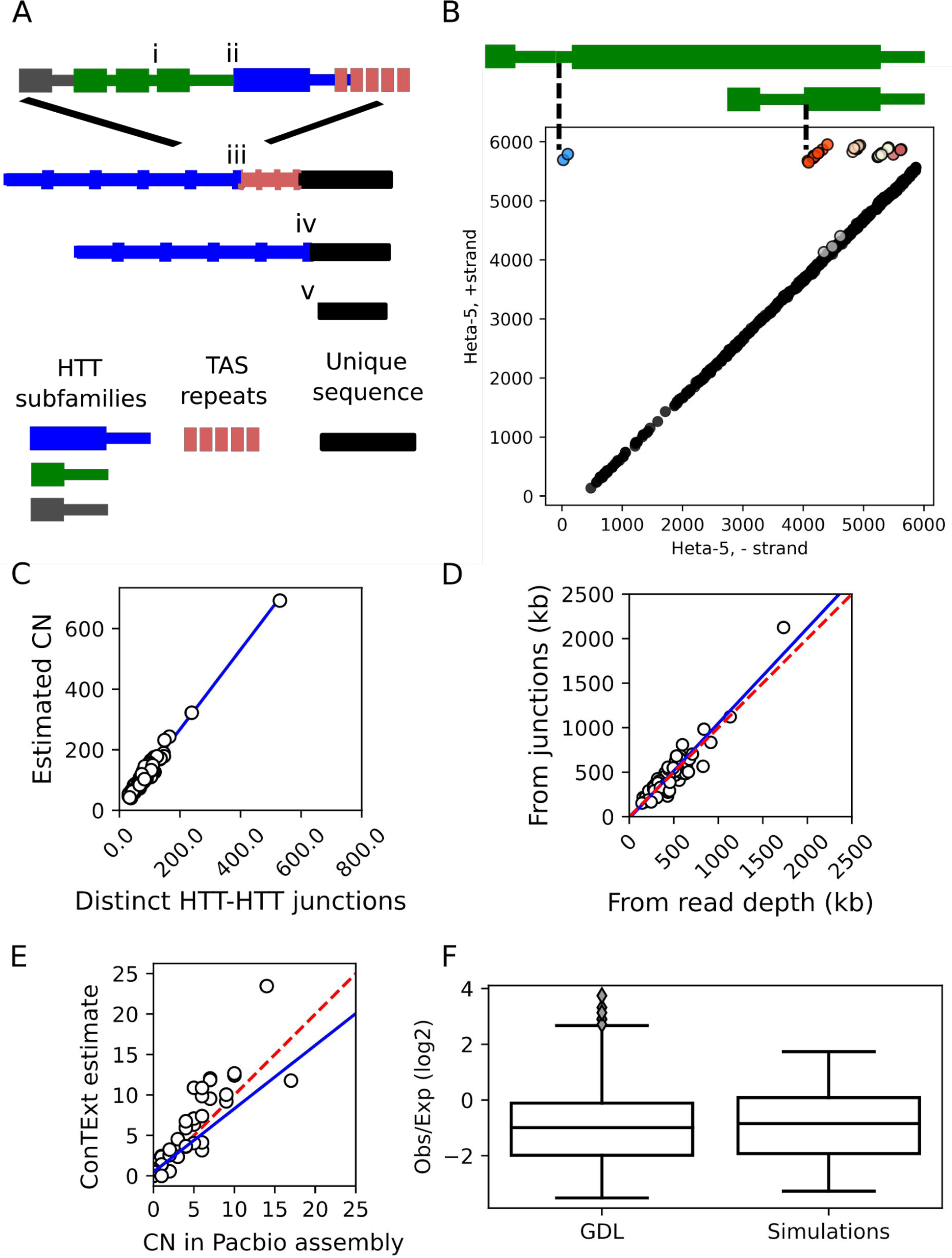
Methodological overview. (A) Graphical representation of a Drosophila telomere and the junctions we use to query telomere variation: i) junctions between adjacent HTTs of the same subfamily, ii) a junction between adjacent insertions of two different HTT subfamilies, iii) a junction between the HTT at the base of a telomere and its neighboring TAS repeat, and iv) a junction between a healed terminal deficiency and unique sequence. The bottom diagram v) depicts the structure of a chromosome with an unhealed deficiency. (B) An example using *HeT-A5* to depict the correspondence between the junctions of adjacent insertions of the same subfamily (i in A) and Illumina read pairs from strain N16. Each dot corresponds to a read pair where both ends map unambiguously to the *HeT-A5* consensus sequence. The Y-axis corresponds to the *HeT-A5* plus strand and the X-axis to the minus strand. The diagonal of dots colored black corresponds to concordant reads that align as would be expected given the consensus element. Non-concordant reads aligning off this diagonal reflect junctions between tandem elements (above the diagonal) or internal deletions (below the diagonal). Small indels may shift reads just above or below the diagonal, for example the grey cluster near the diagonal. Junctions between different families of repeats are detected by considering plots where the X- and Y-axes correspond to different consensus sequences. ConTExt identifies junctions by clustering the non-concordant reads, cluster assignments are reflected by the color of the dots. The five clusters forming a horizontal line across the top of the plot correspond to five distinct tandem junctions between the 3’ end of a *HeT-A5* element and the generally truncated 5’-end of an adjacent *HeT-A5*. Two of the tandem arrangements are illustrated above the plot. (C-F) Comparisons of different approaches for estimating telomere length and HTT copy number, and comparisons against simulated data. The blue lines indicate OLS linear regressions and the red dotted lines indicate a one-to-one relationship for comparison. (C) The relationship between the total number of distinct HTT-HTT junctions identified by ConTExt in each strain (X-axis) and the total HTT copy number inferred from the read depth over these junctions (Y-axis). D) A comparison of total telomere length in each strain estimated from the mapping-quality-filtered read count of junctions (Y-axis) and from coverage of HTT consensus sequences without mapping quality filtering (X-axis). The junction-based telomere-length estimates are obtained by multiplying the inferred copy number of each identified HTT-HTT junction by the length of the distal element inferred from its degree of 5’ truncation. (E) Correlation in CN estimates from simulated Illumina datasets with true copy number. Each dot represents the copy number of an HTT family in one of the five PacBio genomes. The X-axis indicates its copy number estimated from the number of 3’-ends detectable in BLAST alignments between the PacBio assembly and the HTT consensus sequences. The Y-axis is the copy number estimated by ConTExt from data simulated from the PacBio genomes using ART. (F) The downward bias in copy number we observe in the true GDL data is recapitulated in the simulations. The Y-axis is the observed read count divided by the read count expected given the junction’s local GC content. The boxplots depict the distribution of this ratio across all identified junctions in the true GDL data and the simulated data.

With the data organized in this manner, we first determined sequence coverage relative to each HTT consensus sequence, which provides an estimate of the number of nucleotides that each HTT family contributes to the genome. When we use this estimate to calculate total telomere length, we include all reads mapping to HTTs, but when we make statements about the copy number of particular HTT families or subfamilies, we filter out any ambiguously aligned reads to avoid bias from mismapping between related families/subfamilies (see Materials and Methods for details). While this estimates telomere length, the propensity of HTTs to be 5’-truncated means that it is not a direct measure of copy number: coverage of the 3’ end reflects reads derived from all copies, but coverage of the 5’ end reflects only full-length insertions (which may, though, have internal deletions). Second, to more directly estimate copy number, we counted the junctions between adjacent HTT insertions identified by ConTExt. This allows us to infer the number of insertions, the degree to which individual insertions are truncated, and which sequences neighbor a given HTT. To account for the possibility that some junctions are present in multiple copies, we employed partial pooling to estimate copy number from the read counts at junctions. The read depth of the junctions may be downwardly biased by several sources, particularly by mapping quality filtering and the presence of additional sequence in the junction, such as poly-A tails, which we account for by calibrating against a set of HTT junctions with known copy number (see Material and Methods). The copy number estimates from modelling read depth are ∼30% higher than from simply counting junctions, suggesting some junctions are multicopy, but the estimates are highly correlated with each other (slope = 1.3, r = 0.99, Figure 1C). Similarly, estimating the total telomere length from the inferred copy number of each junction and the estimated extent of truncation of the corresponding HTT concords well with the estimates of telomere length computed from raw coverage (slope = 1.06, r = 0.93, Figure 1D). This suggests that our read count model reasonably accounts for the impacts of filtering out ambiguous alignments. Finally, we examine the sequence of aligned reads themselves to assess for variant alleles.

The strains in the GDL were sequenced at relatively low coverage, with the expected read count over individual junctions ranging from ∼10x to 20x across strains. Variation in read depth means that, in some strains, we may be more likely to miss true HTT junctions, but it explains a relatively small fraction of the variation in HTT copy number estimates (Figure S1A). Therefore, to assess how well ConTExt summarizes telomere composition from low coverage NGS data, we used ART to simulate NGS libraries from five available PacBio GDL genomes (Long *et al*. 2018), matching the read depth and insert size distributions to the corresponding strain’s NGS library (Huang *et al*. 2012). We next ran ConTExt on the simulated data and estimated the copy number of the HTT subfamilies using the methods we applied to the true GDL data. We then compared the copy number estimates for each HTT subfamily in the simulated data against the true number evident in the corresponding PacBio assembly. We found these to be strongly concordant, with no evidence of strong upward or downward biases (slope = 1.15, r^2^ = 0.88, p = 1e-16, Figure 1E). We do note one outlier (Beijing line B59) where ConTExt underestimates *HeT-A* copy number, which likely reflects that, despite performing generally quite well, highly divergent copies of a subfamily are more difficult to capture under our alignment parameters. Finally, the original study of SNPs in the GDL reported some residual heterozygosity, which would both double the number of HTT-HTT junctions and halve their read depth in heterozygous tracts. In the real NGS data, we noted that the read depth over HTT-HTT junctions was about half of what was expected given the coverage of unique sequence, which we corrected for by calibrating against HTT junctions with known copy number, i.e. healed terminal deficiencies (see Materials and Methods, “*Correcting for additional biases in read counts over junctions’*). We found that the simulations recapitulate this feature of the data (Figure 1F), suggesting that the alignment process and subsequent filtering of ambiguous reads is likely responsible for reduced coverage over HTT-HTT junctions, rather than residual heterozygosity or underrepresentation of repetitive sequences in the data. We note, however, that our analysis of terminal deficiencies provides evidence of some residual heterozygosity at the telomeres, which we discuss further below. Overall, we find that ConTExt estimates are sufficiently reliable to draw general conclusions about telomere structure in the NGS GDL data.

A final aspect of the dataset that must be considered is that the GDL stocks were collected from five populations and inbred for more than 5 years, meaning that lab evolution may be substantial and heterogeneous across the lines. This could intensify background effects on TE activity, as any polymorphic TE insertions and host alleles that affect their activity will frequently become fixed together in a line, whereas in natural populations they would typically be separated by recombination and independent assortment. In most of our analyses, this should not greatly impact our conclusions. For example, the arrangement of HTTs within telomeres should not be highly sensitive to such continued evolution. Lab evolution, furthermore, may present an opportunity to observe more extreme patterns of some biases that do occur in nature through continued mis-regulation. As we describe below, the clearest impact of continued evolution is evident in our analyses as strain-specific copy-number expansions of some TE families, patterns that are unlikely to be observed in recently collected flies.

### Telomeres are highly dynamic in Drosophila melanogaster

*Drosophila melanogaster* telomeres are comprised of three non-LTR retrotransposon families *HeT-A, TAHRE*, and *TART*, which in our analyses are further subdivided into five *HeT-A* subfamilies (*HeT-A, HeT-A1, HeT-A-2, HeT-A3, HeT-A5*) and three *TART* subfamilies (*TART-A, TART-B1, TART-C*) (see Materials and Methods for details, Table S1 for reference insertion coordinates and Supplementary File 1 for consensus sequences). For clarity, we use the terms “*HeT-A* family” and “*TART* family” when referring to the families. Telomere length has previously been reported to vary by two orders of magnitude (Wei *et al*. 2017). However, that survey was based on qPCR estimates of abundance using a primer set that was designed to detect a single consensus *HeT-A*, in a single North American population, so the absolute size range of telomeres in *D. melanogaster* remains unknown.

To gain a more comprehensive picture of the length and composition of *D. melanogaster* telomeres, we consider how the size and composition of telomeres vary among strains in terms of the sequence abundance and copy number of all HTT families. The median total telomere length in the GDL is 400 kb with a median of 98 HTT insertions per genome, corresponding roughly to 50 kb and 12 insertions per each of the eight telomeres (Figure 2A, Table S2). The majority of insertions belong to the *HeT-A* family (64%), with *TAHRE* (17%) and the *TART* family (19%) contributing fewer insertions (Figure 2B). While these proportions vary across strains, insertions of each of the three families are identifiable in each of the 85 GDL strains. The relative abundance of *HeT-A* and *TART* family elements is roughly equivalent to previous estimates based on Sanger sequencing (Abad *et al*. 2004; George *et al*. 2006). The approximately equal abundance of *TAHRE* and *TART* family elements contrasts with an analysis of the 4R telomere in the reference strain, which contains 8 *TART* family insertions but no *TAHRE* elements (George *et al*. 2006). This may reflect a peculiar structure of the 4R telomere, because analysis of three other telomeres in the reference strain identified 4 *TAHRE* elements (Abad *et al*. 2004). Other strains also show *TAHRE* on a subset of telomeres (Shpiz *et al*. 2007). The majority of HTT insertions are incomplete, with about 80% of *HeT-A* family and *TAHRE* insertions and 93% of *TART* family insertions being truncated to various degrees (Figure 2C). This is similar to what George et al. (2006) observed on telomere 4R, where 75% of *TART* family insertions (6/8) and 66% of *HeT-A* family insertions (10/15) were incomplete. While the proportion of truncated elements varies considerably among strains, we found that there are no consistent differences among populations.

**Figure 2.**
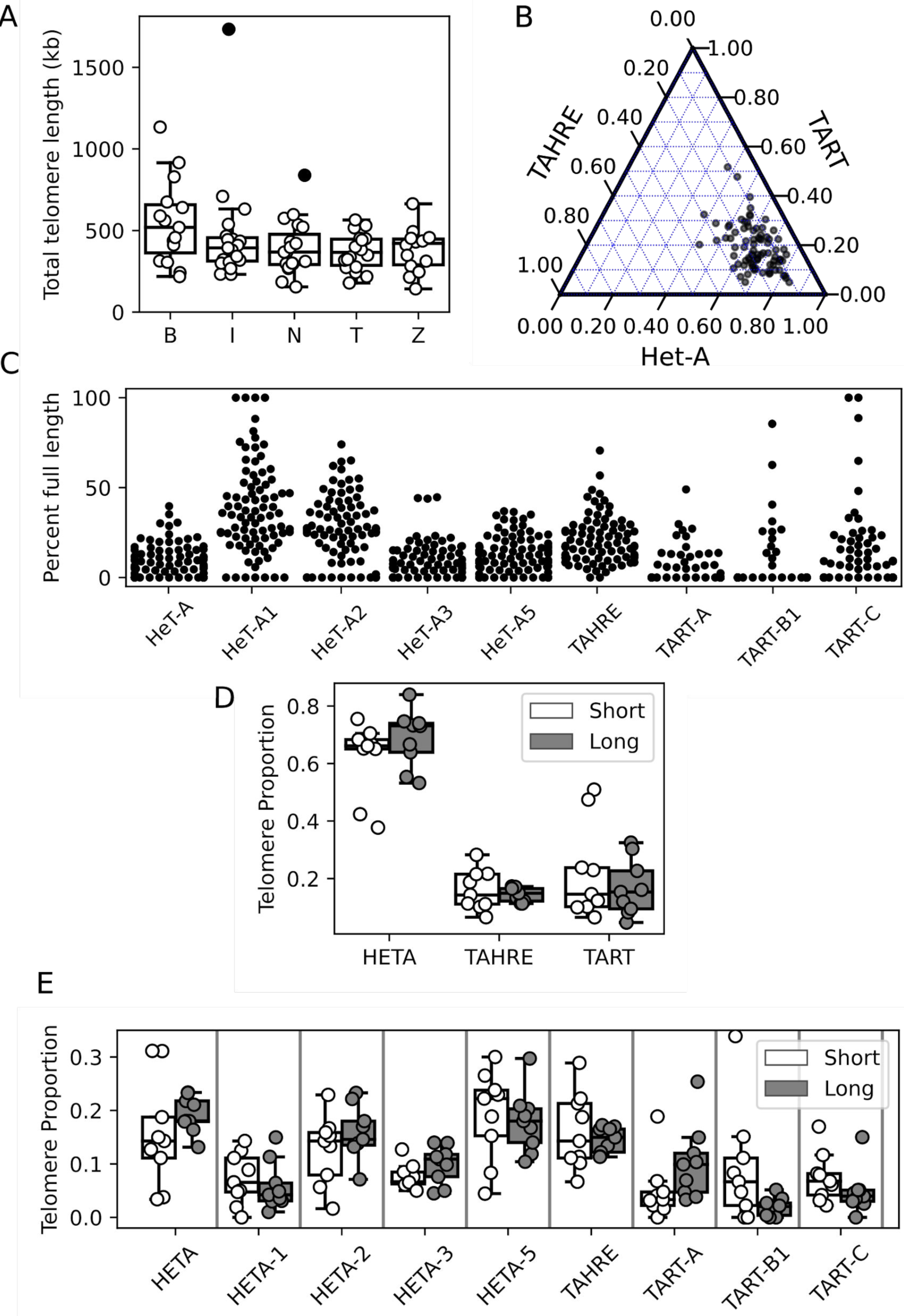
Telomeres are highly dynamic in *Drosophila melanogaster*. (A) Telomere length distribution (in kb) as estimated from HTT sequence abundance for each strain, grouped by population. Filled circles represent outlier strains. B: Beijing, I: Ithaca, N: Netherlands, T: Tasmania, Z: Zimbabwe. (B) A ternary plot depicting the proportion of each HTT family in each GDL strain. The angle of the tick on each axis indicates the corresponding gridline for that HTT family. (C) Proportion of full-length elements per subfamily per strain. (D) Telomere composition depicted by proportion of total telomere length per HTT subfamily as estimated from copy number. White corresponds to short telomere strains (bottom 10%), grey to long telomere strains (top 10%). (E) Telomere composition depicted by proportion of total telomere length per HTT family as estimated from copy number. White and grey are the same as in (D).

Telomere length displays appreciable variation among individuals, with an interquartile range of 193kb and 58 HTT insertions (Table S2). In addition, there is clear potential for extreme gains and losses of telomere sequences, which are observed as significant outliers. Several strains have much shorter total telomere length, with the shortest (ZW140) being 143 kb and having 42 insertions (average of 17 kb or 5 insertions per telomere). Some of the shortest strains are among those that contain unhealed telomere deficiencies (see “Terminal deficiencies are common” below). At the opposite extreme, in I01 the telomeres have expanded to 1.7 Mb of sequence, corresponding to 692 insertions (average of 212 kb or 86 insertions per telomere). The range of telomere lengths is similar to that found in an earlier study of several strains (including the long-telomere strain *GIII*) that used a different approach (Southern blotting) (George *et al*. 2006). There is some population structure to telomere length, with the estimate of mean telomere length in the Beijing population being 110-140kb longer than the other four populations (Ithaca, Netherlands, Tasmania and Zimbabwe), corresponding to 17-40 more insertions per strain on average. This holds after accounting for how differences in read depth across strains impact the probability of not observing a junction that is truly present (Figure S1A). Increases in the number of *HeT-A* family elements drives this heightened copy number in Beijing strains, especially the *HeT-A*1, *HeT-A*2, and *HeT-A*5 subfamilies (Figure S1 B-J).

### HTT family composition is similar in long and short telomere strains

We described above that the Beijing population has elongated telomeres, and that this is driven by expansions of particular *HeT-A* subfamilies. We thus wondered whether telomere elongation reflects escape from host regulation by specific HTT subfamilies. If a specific HTT escapes host control and drives telomere elongation across lines, this subfamily should be over-represented in long telomere strains compared to short telomere strains. We analyzed telomere composition by comparing HTT family and subfamily copy number and relative proportions between long (90^th^ percentile, > 163 HTT insertions) and short (10^th^ percentile, < 54 insertions) telomere strains. At the family level, there is no apparent difference in telomere composition between short and long telomere strains (Figure 2D). As the copy number of *HeT-A* family, *TAHRE* and *TART* family elements increases in long telomere strains, so does their relative proportion, resulting in similar telomere composition. However, some differences arise at the subfamily level (Figure 2E). In long telomere strains, the copy number of *HeTA-1, HeTA-5, TART-B1* and *TART-C* increases less substantially than other sub-families, resulting in a reduced representation in long telomere strains. Altogether, these observations suggest that while telomere length is under host control, HTT sub-families respond differently to a more or less transposition-permissive state, and that long telomere strains do not result from host regulation escape of specific HTT subfamilies.

### Copy number expansions are not specific to HTTs

The pronounced copy number expansions of HTTs we observed in some strains raises the question of whether this is specific to the HTTs. We found that large TE expansions are common in GDL strains for both HTTs and non-telomeric TEs (Figure 3A). Interestingly, in line I01, which has the greatest number of HTTs, *Copia* and *MDG3* (both LTR retrotransposons) have also considerably increased in copy number (Figure 3A, arrow heads; Table S2). Total TE copy number is also highest in strain I01, but across all lines there is no strong general tendency for strains with non-HTT copy number expansions to have longer telomeres. Three non-HTT TE families, *Gypsy1, Zam*, and *Gypsy*, that were defined as inactive under our criteria (see Materials and Methods) are present in few copies in most strains, averaging 2, 3, and 4 copies, but expanded to 20, 37, and 78 copies, respectively, in single strains. For the most part, the copy number expansions all occurred in different strains, and we observe no general relationship between the occurrence of TE expansions and HTT copy number, with the exception of the aforementioned strain I01. It is noteworthy that in this strain, all of the *HeT-A* subfamilies and *TAHRE* have expanded considerably in copy number, but not the *TART* family elements, suggesting differential responsiveness among the HTTs to whatever caused the expansion. The correlation between *HeT-A* elements and *TAHRE* is itself consistent with the proposal that *TAHRE* is the autonomous regulator of *HeT-A* (Abad et al. 2004), such that both families are coregulated. We suspect that most or all of the expansions observed occurred during propagation in the laboratory, because in nature the new insertions resulting from such an expansion would be at low population frequency and rapidly become unlinked from each other due to independent assortment and recombination. In contrast, new insertions can more easily reach high population frequency in the laboratory due to either the small population size or the fixation of permissive variants in piRNA clusters or piRNA pathway genes. We conclude that like any other active and selfish TE, HTTs are primed to take over given the opportunity.

**Figure 3.**
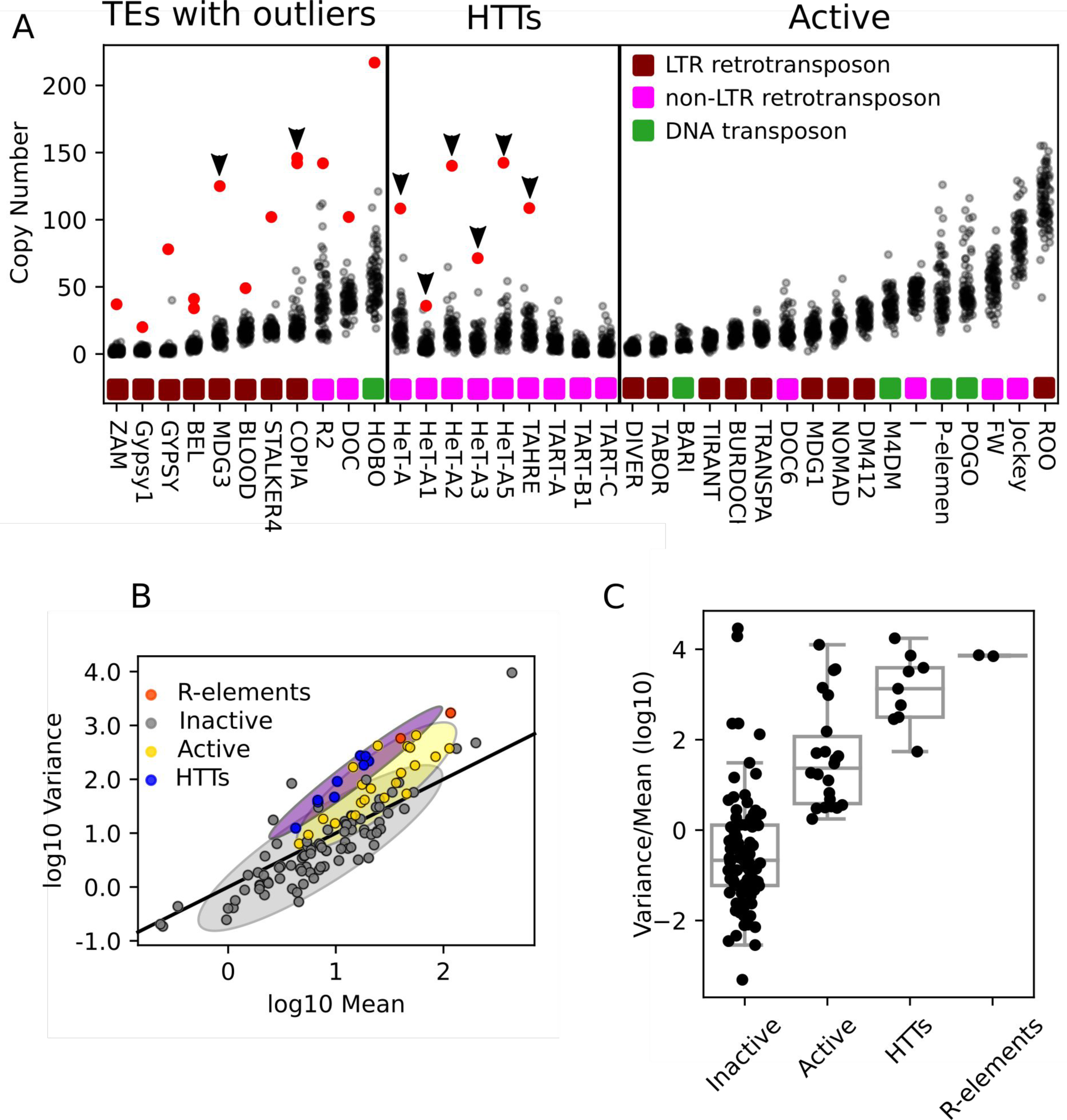
Comparing HTT variability to other TEs within *Drosophila melanogaster*. (A) Copy number of selected TEs per strain as estimated from junctions. Each dot indicates the copy number of a TE family in a single strain, and red dots indicate strains with extreme copy-number expansions (four standard deviations greater than the mean). TEs are grouped as: Left, TEs with copy number outliers; middle, HTTs; right, all other active TEs. Outliers occurring in strain I01 are indicated with arrows. (B) Scatter plot showing the relationship between mean copy number as estimated from junction data of TE families (log scale) and their variance (log scale) across the GDL. Designations of active and inactive TEs are from prior estimates of sequence divergence and population frequency as described in the Materials and Methods. Solid line represents the expected relationship under the assumption of little variation in population frequency and low linkage disequilibrium among insertions. Shaded regions summarize the distributions of mean and variance for inactive (grey), active (yellow), and the HTT and *R*-elements (purple) TE families, covering two standard deviations of bivariate Gaussians matched to the moments of the data they are approximating. (C) Boxplots depicting the distribution of variance-to-mean ratios (log_10_) in each of the four categories of the TEs.

### HTT copy number variation reflect its highly dynamic environment

Given that HTT abundances are highly dynamic across the GDL panel, we wanted to assess whether the extent of this variation was typical of active transposable elements or instead if the HTTs were more (or less) variable in copy number than other TE families. Active elements provide a reasonable baseline because they are in conflict with the genome via transposition. However, the tandem nature of the telomeres additionally permits copy number expansions and contractions through unequal exchange, which may heighten the variability in copy number. Two other TE families, the *R1* and *R2* elements, are also tandemly arrayed within the multi-copy ribosomal RNA genes (rDNA) and thus reside in a dynamic environment. The *R1* and *R2* elements therefore provide a comparison of the HTTs with TEs whose copy number evolves both by transposition and unequal exchange but are not known to provide an essential function.

We sought to leverage population genetic theory for the relationship between the mean and variance of TE copy number distributions to interpret the empirical relationship we observed in the GDL. Charlesworth and Charlesworth (1983) highlight that the variance in TE copy number should depend only on the mean and variance of the allele frequency spectrum of TE insertions and the degree of correlation in the presence/absence of distinct insertions (linkage disequilibrium). The intuition is that whatever dynamics and processes underlie the copy number evolution of a TE family, each occupiable locus contains an insertion in some fraction of individuals in a population. The set of loci containing insertions in an individual genome is sampled from this larger population of insertions, each insertion allele being present with probability equal to its population frequency. Assuming that the number of *de novo* insertions per generation is small relative to the number of preexisting insertions, the copy number in an individual genome is then the sum of Bernoulli trials with potentially different success probabilities. If these trials are independent and have identical population frequencies, TE copy number will be Poisson distributed and the variance of copy number across genomes will be equal to the mean genomic copy number. If they are independent but the population frequencies vary, this can be described with a Poisson-binomial distribution, and the variance will be less than the mean. In reality the presence/absence of insertions may not be independent of each other (termed “linkage disequilibrium”) due to a variety of factors, including genetic linkage, selection, and demographic history and population structure. This correlation in the presence/absence of insertions will increase the variance of TE copy number. These factors are synthesized in the mean-variance relation

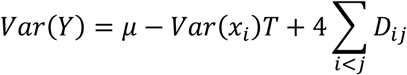

where *μ* is the population mean copy number, *T* is the number of occupiable loci, *Var*(*x*_*i*_) is the variance in population frequency of TE insertions and *D*_*ij*_ is the linkage disequilibrium of insertions (Charlesworth and Charlesworth 1983). The principal insight is that TE families whose insertions vary over a range of population frequencies should display decreased copy number variation. On the contrary, those families whose insertions are in linkage disequilibrium should display elevated copy number variation. This is what we expect for HTTs, since their tight clustering at eight telomeric tandem arrays may lead to co-transmission or loss of many insertions at once.

We examined the mean-variance relationship of TE copy number across families, considering two different estimates of TE copy number (see Materials and Methods) which gave similar results. We found that inactive TE families are generally less variable in copy number than expected relative to a Poisson expectation (Figures 3B, C and S2 A,B). This is consistent with theory, as relatively older (but not ancient) insertions should have more variation in their population frequencies than recent insertions. This is because older insertions have had more time to change in population frequency, leading to a less variable copy number distribution (due to *Var*(*x*_*i*_)*T* > *0*). In contrast, active TEs not only show greater copy number variation than inactive TEs, but are generally more variable with respect to the Poisson expectation. This greater-than-expected variability may reflect linkage disequilibrium or population structure during the TE family’s invasion. We consider continued copy number evolution in the lab stocks (itself a form of population structure) a likely explanation. However, compared to other active elements, HTT copy number is even more variable (Figure 3B, C, B and S2A, B), which could reflect instability of the genomic regions they occupy, coupled with high linkage disequilibrium (*4* ∑_*i*<*j*_ *D*_*ij*_ > 0).

That is, while other TE families are dispersed across the genome, all HTT insertions are clustered at eight tandem arrays, such that many insertions will be transmitted or lost together. Notably, the *R*-elements also display elevated variability comparable to the HTTs (Figure 3B, C and S2 A,B). We conclude that both the HTTs and *R*-elements are more variable in copy number than other active TEs. This could reflect that these elements are more prone to continued CN evolution during stock maintenance, but it is striking that it is those TE families known to be organized in tandem arrays that are the most variable. We suggest therefore that the heightened copy number variation displayed by the HTTs and R-elements reflects both the dynamic nature and the tight physical linkage of the genomic environments caused by the tandem array organization of these families.

### The high sequence diversity of HTTs is driven by active variants

Sequence evolution of TEs determines their activity and their potential to escape from host suppression. HTTs are highly divergent among species, which has been suggested to reflect continued conflict with their host genome (Lee *et al*. 2017; Saint-Léandre *et al*. 2019). We propose that the instability of their genomic niche might also contribute to HTT sequence variation. The heightened copy number variation we observe for the HTTs (and *R*-elements) ought to increase the stochasticity with which lineages go extinct, allowing sequence polymorphism to fix more rapidly in them. We sought, therefore, to understand how the within-species sequence diversity of the HTTs compares to that of other active TEs, especially the rDNA-restricted *R*-elements.

For each TE family under consideration, we estimated the copy number of each of the four possible alleles (A, T, G, C) from the number of reads supporting that allele at a given consensus position in a particular strain (Figure 4A; Materials and Methods). We then used these allele copy number estimates to compute the sequence diversity among insertions of a given family in the GDL. The diversity estimates across all insertions in the GDL concord well with those reported from analysis of the ISO-1 reference genome (Kendall’s tau = 0.59, p = 6.6e-5) (Figure S2C; Kaminker *et al*. 2002). We find that HTT subfamilies are relatively diverse in their sequence composition, though not obviously more so than other active TE families (Figure 4B). Such sequence diversity could reflect heterogeneity among the transposition-competent elements. Alternatively, the active elements might be relatively homogeneous, but distinct in sequence from older inactive insertions that are relics of prior invasions. This alternative scenario applies to TEs such as *I*-element and *Hobo*, which have invaded *D. melanogaster* multiple times and left both recently active and degenerate copies from past invasions (Bucheton *et al*. 1992; Boussy and Itoh 2004). The shape of a phylogeny from complete sequences of all TEs in the GDL would easily distinguish these scenarios (Khan *et al*. 2006). However, we cannot reconstruct individual elements, much less their phylogeny, with short reads. To circumvent this challenge, we developed an approach to leverage our observation that the copy number variation of active TE families tends to be overdispersed and that of inactive families tends to be underdispersed (Figure 3B, C). We reasoned that high diversity at positions where both the major and the most abundant minor alleles display more variable copy number than expected (see previous section) is suggestive of heterogeneity among the active elements of the family. On the other hand, if the major allele is more variable but the minor allele is less variable than expected, this instead suggests that the active elements are homogeneous at that position, with a distinct population of inactive copies that are divergent from the active lineage (Figure 4A). This inference is admittedly indirect, and could be obscured by the difficulties of mapping reads from highly diverse TE families. Our focus on recently active TE families, however, should ameliorate this concern, and the concordance between our sequence diversity estimates for these recently active families and those obtained from prior analysis (Figure S2C) indicates that these are not highly diverse TE families and that our estimates are not strongly biased.

**Figure 4.**
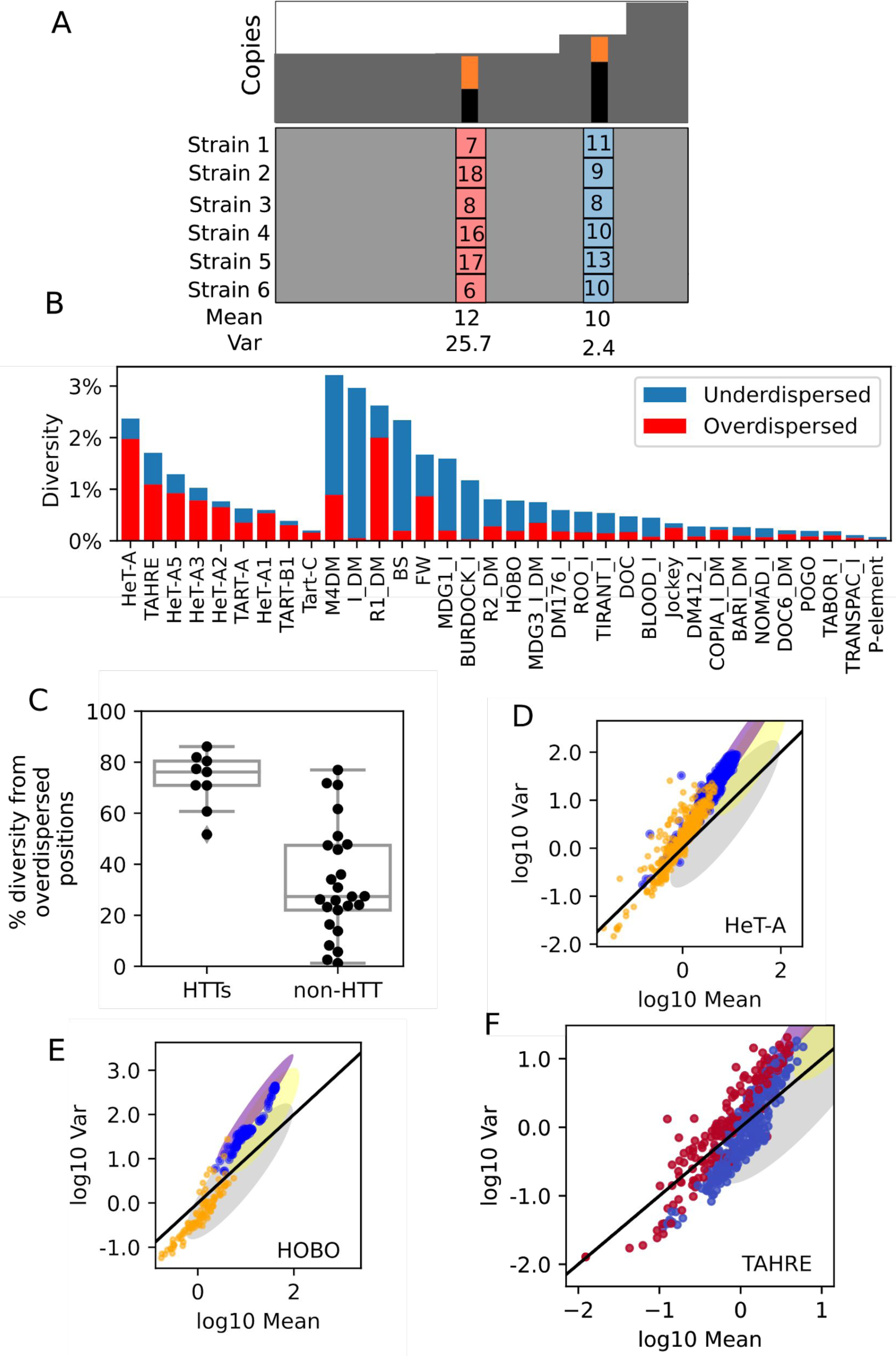
HTT copy number and sequence variation in the context of other TEs. (A) A schematic of how allele copy number is analyzed. Top: Hypothetical copy number of a TE family in a single strain, including two positions where polymorphisms exist such that the copy number can be partitioned into the major (black) and minor alleles (orange). Bottom: A schematic depicting hypothetical copy number variation of the most common minor allele across six strains. If the variance in copy number is greater or less than the mean, the copy number variation is called overdispersed (light red) or underdispersed (blue), respectively. (B) Sequence diversity across all copies of each active TE family in the GDL, estimated from the depth of reads supporting each possible allele. The contribution of positions where the minor allele displays overdispersed copy number variation, suggesting a variant in an active element, is indicated in red. The contribution of positions where the minor allele displays underdispersed copy number variation, suggesting a variant in an inactive element, is indicated in blue. (C) Boxplots depicting the fraction of diversity contributed by positions with overdispersed minor alleles for both HTTs and non-HTT active TEs. The difference in medians is 49% (p = 3e-4, permutation test, 100,000 permutations). (D-E) The mean-variance relationships of *HeT-A* (D) and *hobo* (E) broken down by the copy number of the major and minor alleles. Each dot reflects the observed mean and variance of the copy number of the major (blue) and minor (gold) alleles of positions with >0.1 sequence diversity. For reference, the shaded regions are re-plotted from Figure 3B. (F) The mean-variance relationship of *TAHRE*’s minor alleles, colored by whether the minor allele is found in the heterochromatic *TAHRE* insertions (blue) or is likely telomeric (red). Shaded regions are re-plotted from Figure 3B.

Breaking diversity down into the proportion driven by alleles that vary much in copy number (overdispersed) and those that do not (underdispersed), we find that in HTTs, much of the sequence diversity comes from positions with overdispersed minor alleles, while in most other active TE families the diversity largely corresponds instead to positions with underdispersed minor alleles, suggestive of variants in older insertions. (Figure 4B, C). Consistent with a population of heterogeneous active elements, when we consider the mean-variance relations of the major and minor alleles, we find that in the HTTs both generally display the same mean-variance relationship we observed for HTT copy number and tend to be more variable compared to other TEs, as is seen, for example, in *HeT-A* (Figure 3D, Supplemental File 2). In contrast, for most of the other TE families with high sequence diversity including *Hobo* and *I*-element, the variance of the minor allele tends to be shifted down from that of the major allele and is generally less variable, as is seen in *Hobo* (Figure 4E; similar plots for the other TEs in Figure 4B can be found in Supplemental File 2). We suggest that this pattern of sequence diversity of non-HTTs is largely driven by degenerate inactive copies, with only a small fraction of positions in most elements reflecting divergence among active elements. This is consistent with the known invasion history of *Hobo* and *I*-element (Bucheton *et al*. 1992; Boussy and Itoh 2004). *TAHRE* is one exception to the tendency of the HTTs to show mainly highly variable minor alleles, as it appears to have a subset of less variable minor alleles (Figure 4F), suggesting the presence of inactive copies outside of the telomere. We subsequently confirmed the existence of nearly fixed non-telomeric *TAHRE* elements harboring 214/249 of these underdispersed alleles in unmapped heterochromatin (see “*HTT transposition is restricted to the telomeres*” below), providing additional validation that, despite its indirectness, underdispersion of allele copy number can indeed be indicative of alleles in older insertions. To investigate whether this heightened diversity observed in the HTTs is characteristic of TEs residing within unstable genomic regions, we considered the sequence diversity of the *R*-elements. However, while *R1* also shows high levels of diversity (Figure 4B), the presence of hundreds of tandemly arrayed *R1*-elements per genome evolving independently of the active elements means we cannot distinguish whether this diversity is among the active elements or involves those comprising the tandem array (Roiha and Glover 1981; McGurk and Barbash 2018). A cleaner comparison is with *R2*, which does not form tandem arrays. Unlike the HTTs, *R2* displays considerable variation consistent with inactive and degenerate copies, which likely reflects accretion from the edge of the rDNA array. This accretion of degenerate and fragmented copies of repeat units is commonly observed at the edge of satellite arrays (McAllister and Werren 1999). The absence of accreted HTT relics may simply reflect the action of telomere erosion, which readily deletes telomeric sequence, including sequence accreted into the subtelomere. *R2* displays lower levels of sequence diversity among active copies (Figure 4B, Supplementary File 3), which may indicate that even for TEs within unstable niches, the HTTs are evolving rapidly, but this is a limited comparison because we are comparing to only one element. However, we can more confidently conclude that the high sequence diversity of HTTs is driven by active variants whose appearance and maintenance is favored by the unstable genomic niche they occupy.

### HTT transposition is restricted to the telomeres

While HTT transposition has been apparently restricted to the telomeres for millions of years, the retention of intact endonuclease ORFs in *TART* and *TAHRE* suggests that HTTs could potentially transpose elsewhere in the genome (Casacuberta 2017). Transposition of HTTs outside the telomere might reflect selfish HTT behavior that is in conflict with the host (Saint-Léandre *et al*. 2019). Non-telomeric *HeT-A* family and *TART* sequences are found on the heterochromatic Y chromosome of *D. melanogaster* and other Drosophila (Danilevskaya *et al*. 1993; Losada *et al*. 1997; Agudo *et al*. 1999; Berloco *et al*. 2005). However, Berloco et al. (2005) noted that “these non-telomeric tandem repeats differ from the telomeric elements in at least two respects: they are interspersed with other repetitive sequences and they are not always oriented in the same direction” and postulated that chromosomal rearrangements rather than non-telomeric insertions were responsible. These structural differences suggest that non-telomeric HTT copies may result from gene conversion, capture by other TEs, or rearrangements, rather than by non-telomeric insertion (Danilevskaya *et al*. 1993; Agudo *et al*. 1999).

Outside of the Y chromosome, there are two non-telomeric regions of the mapped *D. melanogaster* reference genome that contain sequence with >80% homology to the HTT consensus sequences, both on 3L: a ∼700 bp stretch between 16,576,994-16,577,693 with ∼95% identity to a fragment of the *TART-A* terminal repeat and a 15 kb stretch between 25,216,958-25,230,409 containing fragments with lower identity (<85%) to *HeT-A, TAHRE*, and *TART-A* sequence. Their short lengths (several hundred base pairs or smaller) and adjacency with other HTT sequence suggest that they either arose from recombination-mediated transfer of telomeric sequence (Agudo et al 1999) or represent ancient insertions that have largely degraded. Non-telomeric insertions were recently reported as common, however, in the *melanogaster* subgroup species *D. rhopaloa* (Saint-Léandre *et al*. 2019). We therefore used ConTExt to search the *D. melanogaster* GDL for HTT insertions in assembled regions as well as in other repeats. We identified 34 junctions between HTT 3’-ends and subtelomeric unique sequence, these reflect healed terminal deficiencies and are discussed separately below. Beyond the subtelomere, we only found junctions supporting seven loci in the mapped chromosomes potentially harboring non-telomeric HTT sequence. Both 3’ and 5’ junctions of the aforementioned *TART-A* terminal repeat fragment at ∼16,576,000 on 2L were identifiable, with at least one of the junctions detectable in 82/85 strains. The only other non-telomeric HTT with both 3’ and 5’ junctions detectable was a short *HeT-A* fragment spanning 5,276 - 5,574 of the consensus at position ∼7,537,400 on 3L in strain I29. The remaining five loci were supported by only a single junction with a *HeT-A* subfamily, and the only one present in more than one strain (3/85) falls at the edge of the previously mentioned stretch of 3L harboring degenerate *HeT-A* fragments (25,229,995). Furthermore, none had both 5’ and intact 3’ junctions; it is therefore challenging to distinguish these loci from sequencing artifacts. Assuming that they are genuine, it is likely that these fragments arrived outside the telomere due to a recombination-based mechanism, though two of the seven non-telomeric HTTs may have intact 3’-ends and thus could represent rare non-telomeric insertions. We conclude that non-telomeric insertions of HTT elements are very rare or negligible in *D. melanogaster*.

For perspective, we searched for *R*-element insertions outside the rDNA. While domestication of the HTTs could keep them constrained to the telomeres, there are unlikely to be similar constraints on the *R*-elements because they serve no obvious function. Rather, their restricted localization to the rDNA is probably a property purely intrinsic to the *R*-element transposition machinery. Thus, the extent to which we detect *R*-element insertions outside of their niche provides both a positive control validating our approach and insight into the extent to which niche specificity can be maintained by TEs alone without external control by the host. We identified no instances of *R2* elements inserted outside of rDNA units in the GDL data. By contrast, we found 33 insertions of *R1* outside of rDNA units where both the intact 3’-end and (often truncated) 5’ junctions were identifiable. Most such junctions were restricted to single strains, though one insertion in the 3L pericentric heterochromatin was present in 70/85 strains, suggesting it occurred more distantly in the past. Overall, while this corresponds to only 0.34% of all *R1* elements in the GDL, it nonetheless reflects more insertion site promiscuity outside of its typical niche than we observe for the HTTs. On the whole, though, the tight restriction of the *R*-elements and HTTs to their respective niches suggests it is largely a property of the elements themselves and that host control may not be required to maintain highly specific insertion sites.

However, while we did not identify clear evidence of non-telomeric HTT insertions in well-mapped regions of chromosomes, several of our observations suggest there may be copies of *TAHRE* outside the telomere but within unmapped heterochromatic regions. First, a subset of *TAHRE* SNPs vary less in copy number than expected of alleles found in active elements, suggesting the possibility of inactive copies outside the telomeres (Figure 4F). Second, we find that few strains lack full-length *TAHRE* insertions in contrast to the other HTTs (Figure 2C), suggesting there may be tandem *TAHRE* copies that are rarely lost. Finally, we observe an *R1* inserted into *TAHRE* sequence which has reached moderate population frequency (12/15) among Beijing lines. Given the instability of the telomeres, we think it unlikely that such structures could be present in so many genomes if they involved telomeric *TAHRE*.

We therefore searched for non-telomeric *TAHRE* elements in unmapped heterochromatic sequences missing from the reference genome. As the unmapped contigs in the reference genome are too short to determine whether *TAHRE*-containing contigs are surrounded by telomeric repeats or instead embedded in non-telomeric repeats, we searched for heterochromatic *TAHRE* insertions in the PacBio long-read assembly (GCA_000778455.1) of the reference strain ISO1 (Berlin *et al*. 2015). We identified one 140kb scaffold (JSAE01000744) which contains three tandem *TAHRE* insertions embedded among many other TEs. These are highly similar to each other, though 5% divergent from the *TAHRE* consensus. Both the 3’- and 5’-ends of two insertions are evident and they form tandem junctions. These *TAHREs* likely correspond to the full-length *TAHRE* that we found above in most strains. A majority (214/249; 86%) of the underdispersed *TAHRE* minor alleles (which led us to search for non-telomeric *TAHRE*s) correspond to polymorphisms present in these heterochromatic elements (Figure 4F), indicating that these are also the cause of the underdispersed alleles and providing validation for our inference that underdispersion reflects older insertions. Some polymorphisms within these non-telomeric copies are present in more than three copies and display overdispersed variation more consistent with active elements, suggesting that some extant active *TAHRE*s are closely related to the lineage that gave rise to these tandem *TAHRE*. Both the *gag* and *Pol* ORFs are interrupted by several nested TE insertions that are present in most (82/85) strains. Tandem TEs indeed can form by multiple, independent insertions at the same locus, but generally constitute a minority of total insertions and often form at strong consensus insertion sites (McGurk and Barbash 2018). If these tandem *TAHRE* elements reflect true non-telomeric insertions, at least two independent *TAHRE* insertions must have occurred at this locus. Given that we observed no single insertions outside of the telomeres, we consider double insertion outside of the telomere to be unlikely (although it is possible that non-telomeric *TAHRE* insertions occurred more frequently in the past). We instead propose that these *TAHRE* elements originated as insertions within heterochromatin-adjacent telomeres, XR or 4L being likely candidates, and were subsequently pushed into the adjacent heterochromatin by accretion from the edge of the array.

### Terminal deficiencies are common

The high degree of copy number variability displayed by the HTTs appears to be shaped by the dynamic nature of the telomere. We sought, therefore, to further characterize the stability of the telomeres. Beyond the action of unequal exchange between telomeres to amplify and delete sequence, there are other possible mechanisms by which telomeric sequence may turn over. First, the end replication problem directly causes telomere erosion, which in some cases may reduce the chromosome ends to the subtelomeric TAS arrays or even into centromere-proximal unique sequence. Second, DNA double-strand breaks in the subtelomere may result in sudden terminal deficiencies.

We consider three lines of evidence to identify deficiencies. First, to identify the subset of terminal deficiencies that extend through the HTT array and subtelomeric repeats into unique sequence, we searched for a sharp loss of (homozygous) or reduction in (heterozygous) read depth over telomere-proximal unique sequence. Second, deficiencies which have been healed by new HTT insertions can be detected as junctions between the 3’-ends of HTT elements and subtelomeric unique sequence (Biessmann *et al*. 1990b). Such insertions onto the end of a broken chromosome can be readily distinguished from chromosome-internal insertions, which would produce junctions with unique sequence involving both the 3’ and 5’ ends of the HTT. Considering any subtelomere that met either of these criteria, we found evidence for 56 terminal deficiencies in 48/85 strains (Figure 5A, Supplemental File 3). As a third and independent test, we looked for concomitant deletion of the subtelomeric satellite. The left arms of chromosomes 2 and 3 harbor arrays of the same subtelomeric satellite, TAS-L. If we have identified true terminal deficiencies of 3L, the TAS array should be lost from those chromosomes and TAS-L copy number reduced. Consistent with true terminal deficiencies on 3L, the TAS-L copy number is roughly half that of strains without detectable deficiencies in strains with HTT-3L junctions (Figure 5B).

**Figure 5.**
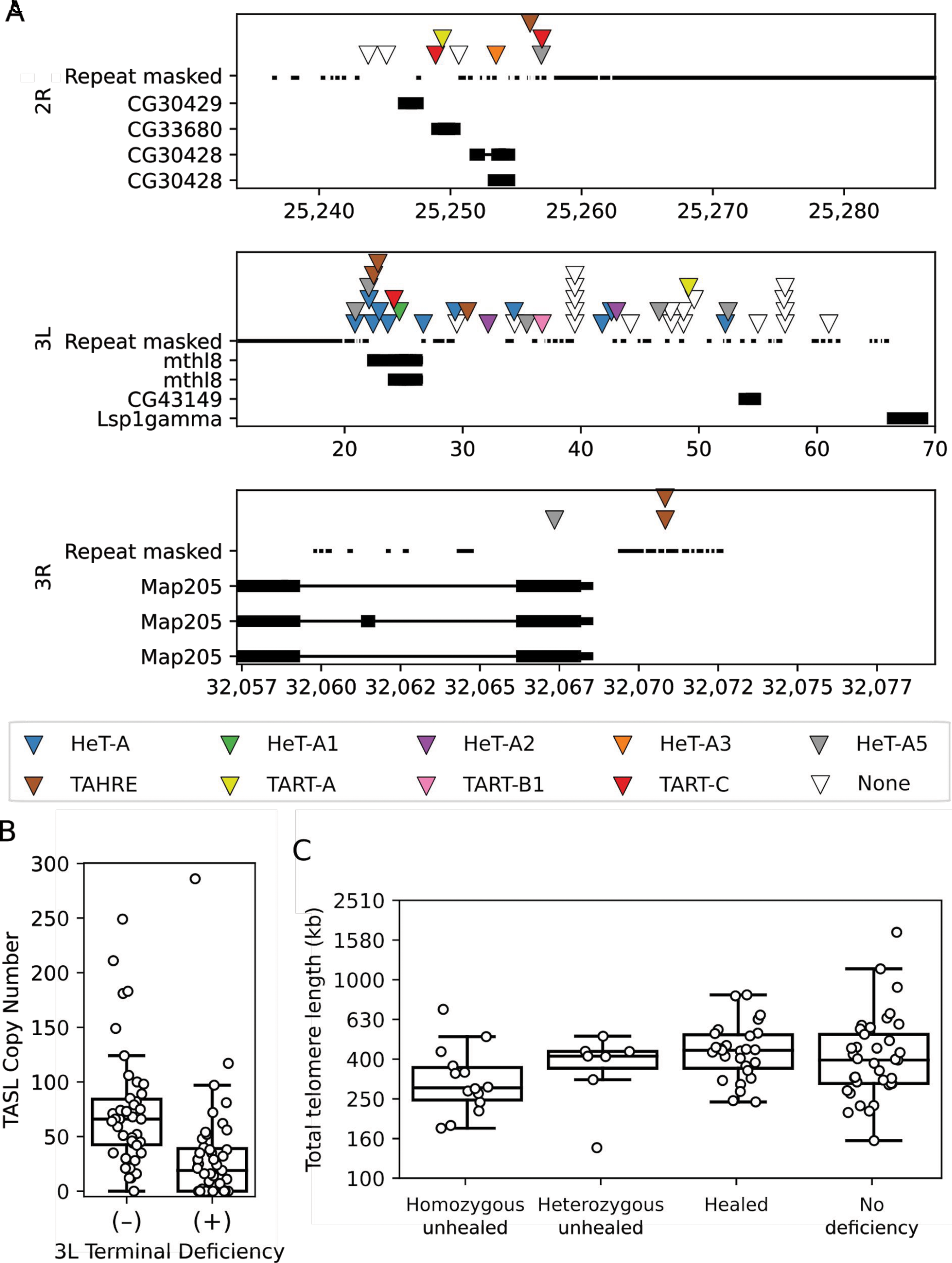
Terminal deficiencies are frequent and tend to be healed by *TART* family elements. (A) Location of all identified terminal deficiencies (triangles represent individual terminal deletions) and elements involved (see colour legend; “none” indicates a terminal deficiency lacking any HTT-subtelomere junction) across all lines. Thin and thick black bars represent UTRs and exons respectively, thin lines represent introns. Top, Chromosome 2R; middle, Chromosome 3L; bottom, Chromosome 3R. X-axis scales are in kilobases; note that the telomeres are to the right for 2R and 3R, and to the left for 3L. (B) TAS-L copy-number boxplot in strains with (+) or without (−) deficiencies on 3L. (C) Boxplots comparing the total telomere length of strains with homozygous deficiencies and heterozygous deficiencies without observed HTT-subtelomere junctions (unhealed) to the telomere length of strains with healed deficiencies and without deficiencies. Strains with healed deficiencies (whether homozygous or heterozygous) will have eight telomeres whereas those with homozygous unhealed deficiencies will have only seven. Note that the Y-axis is in log_10_-scale.

Of the 56 terminal deficiencies, 14 display evidence of heterozygosity, suggesting these are not fixed in the fly stocks and perhaps arose after collection. The remaining 42/56 deficiencies are all consistent with homozygous deficiencies, as expected in inbred strains. Of the 56 deficiencies, 34 showed evidence of an HTT-subtelomere junction, indicating that the lost telomere has been replaced by a new HTT array. All but five of those 34 deficiencies extend sufficiently far into unique sequence that we could confirm a clear drop in coverage after the HTT-subtelomere junction. For 22/56 deficiencies, however, we observe only a loss of subtelomeric unique sequence based on read depth but no evidence of repair in the form of an HTT junction. Some of these may reflect our failure to detect a junction that is truly present, for example due to variation in read counts or repeat-masked sequence (note that five putatively unhealed deficiencies sit at the edge of masked sequence, Figure 5A middle panel). However, total telomere length is generally lower in these strains with putatively unhealed deficiencies compared to strains where a HTT junction is identifiable (Figure 5C), consistent with truly missing a HTT-based telomere. This suggests that a remarkably high proportion of telomere deficiencies are not healed by HTTs.

At least 39 of the deficiency breakpoints are independently derived, involving different sequence coordinates (Figure S2D). Most (44/56) are found on 3L and often delete the most distal gene, *mthl8*, with a few also deleting the next annotated gene *CG43149*; a similar high rate of 3L terminal deficiencies was discovered by Kern and Begun (2008). We note that the density of deficiencies is twice as high distal to *mthl8* than it is between *mthl8* and *CG43149*, suggesting that deletions of *mthl8* nonetheless have some deleterious effect even if not as serious as loss of *CG43149* (Figure 5A, middle panel). Only 9 and 3 deficiencies are present on 2R and 3R, respectively, but the distribution of these are also limited by genes within several kilobases of the telomere, with few extending beyond *CG30429* on 2R and beyond *Map2O5* on 3R. We observed no deficiencies on 2L, likely because there is only 400 bp of non-repetitive sequence between the gene *lethal (2) giant larvae* and the TAS array. Mutations in *l(2)gl* are homozygous lethal; therefore terminal deletions removing *l(2)gl* would not be expected in the GDL lines we sampled because they have been inbred to homozygosity, but they are common on heterozygous chromosomes in natural populations and in heterozygous lab stocks (Golubovsky 1978; Green and Shepherd 1979; Mechler *et al*. 1985; Roegiers *et al*. 2009). Taken together, our analysis reveals that terminal deletions occur at high frequency on all chromosomes, and also highlights the fitness impacts of terminal erosion and deletion.

### Patterns of HTT insertions suggest different families are active under different conditions

The presence of distinct families and divergent subfamilies of TEs is a common feature of Drosophila telomeres and opens up the potential for specialization of elements and differential regulation by the host genome. We investigated two possible cases. First, we hypothesized that HTT specialization could affect patterns of HTT interspersion. For example, competition among elements that is resolved by limiting their expression to different cell types or development stages would result in finding adjacent copies less frequently than expected. Conversely, cooperation among elements that requires their co-expression for successful transposition would result in an increased frequency of interspersed insertions. We found that elements of the same subfamily tend to neighbor each other, suggesting that they tend to be reverse transcribed multiple times at a given chromosome end (Figure 6A, B). We also observed that elements of the *TAHRE* and *HeT-A* families are generally interspersed as would be expected under random ordering. *TART* family elements, however, are less commonly found adjacent to *HeT-A* family and *TAHRE* insertions than expected. Further, the tendency of *TART* family elements to be found adjacent to their own subfamily is much stronger than that of *HeT-A* subfamilies and *TAHRE*. Clustering of *TART* family elements has been previously detected (George *et al*. 2006). The interspersion of *HeT-A* family and *TAHRE* elements contrasted with the low frequency of junctions of either of them with *TART* family elements may be a consequence of the *HeT-A* family relying upon the expression of *TAHRE*’s polymerase for reverse transcription, implying that the *HeT-A* family and *TAHRE* mobilize together (Shpiz *et al*. 2007). It further suggests that *TAHRE* and the *TART* family may not be active at the same time. To confirm these observations, we examined the assembled telomeres in the release 6 reference genome (Hoskins *et al*. 2015), and observed several instances where multiple full-length copies of the same element are in tandem (Figure 6C). We conclude that the pattern of HTT arrays are not random, and that this likely reflects the dependence of the *HeT-A* family on *TAHRE’*s retrotransposition activity as well as possible specialization of the *TART* family versus *TAHRE*. The three *TART-A* elements in tandem on the X chromosome telomere, however, share terminal repeats, which is consistent with recombination between the 3’ and 5’ terminal repeats (Ke and Voytas 1997) and the principal mechanism by which TEs with terminal repeats generate tandems in Drosophila (McGurk and Barbash 2018). Thus the higher rate of self-adjacency observed for *TART* family elements may reflect recombination between termini as previously suggested by George et al. (2006).

**Figure 6.**
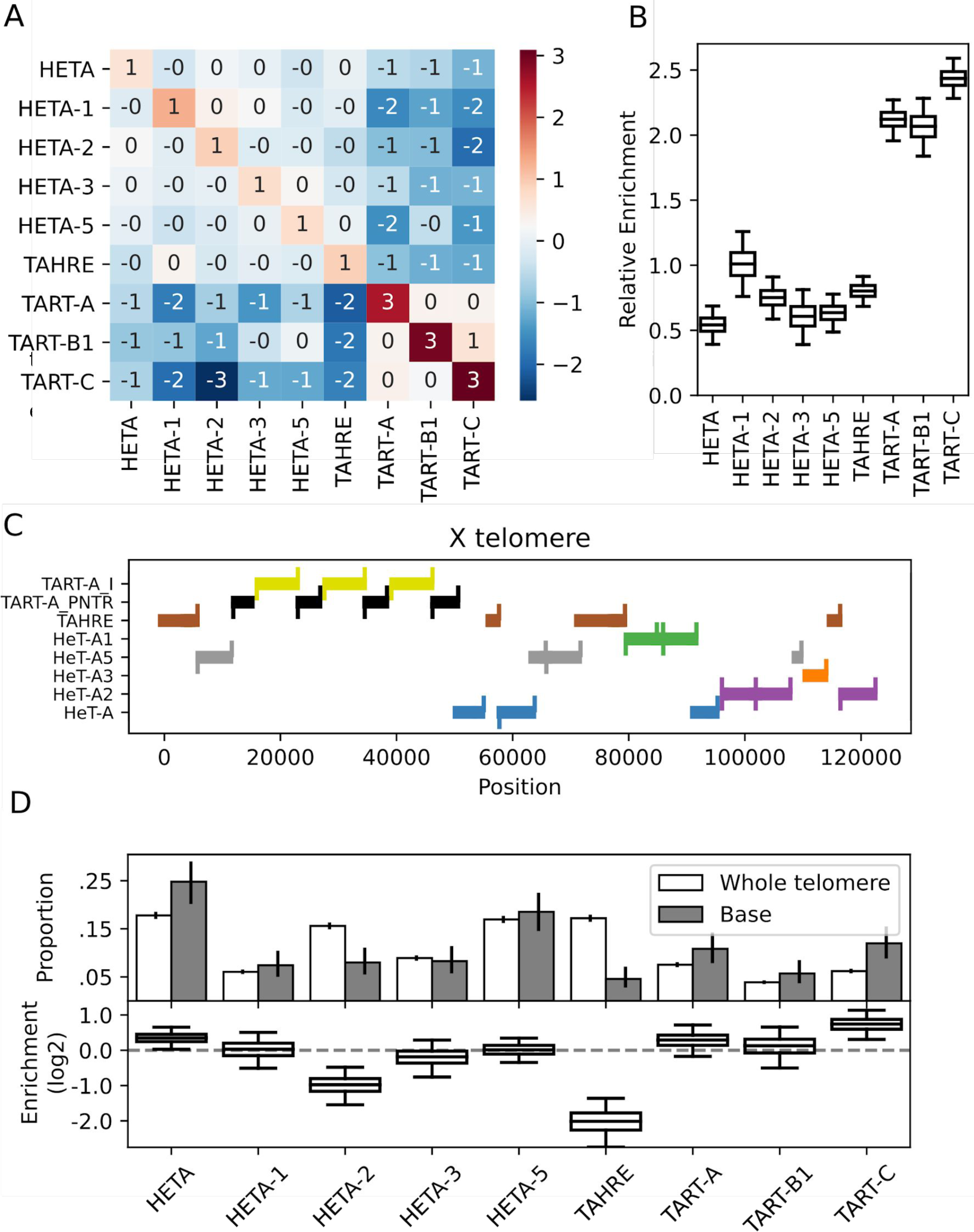
HTT insertions tend to be enriched adjacent to themselves. (A) The observed frequency with which two HTTs neighbor each other relative to the expected frequency (log_2_). (B) Boxplots of the posterior distributions describing the degree to which elements tend to neighbor themselves (log_2_). The whiskers reflect the 95% credible intervals. (C) A visualization of the HTT subfamilies, depicted as thick bars, in the X chromosome telomere of the Release 6 reference. We depict alignments with at least 90% identity to the consensus; if a region is homologous to two elements, we assign it to the element with the greatest homology, which was only an issue due to some homology between *TART-A* and *TART-C*. The upward ticks indicate the 3’-end of an element and the downward ticks the 5’-end; a full length insertion has both ticks. TART-A_PNTR is the TART-A Perfect Near-Terminal Repeat. (D) Top: The observed proportion with which each subfamily is found anywhere in the telomere (white) or is the first HTT found at the base of a telomere (grey). The error bars are 95% credible intervals computed analytically for Dirichlet-Multinomial models with uniform priors. Bottom: Boxplots summarizing posterior samples of the relative enrichment (log_2_) of each subfamily at the base of the telomere, accounting for telomere composition differences across strains. The whiskers span the 95% credible interval, determined as quantiles of the posterior sample.

We also considered the possibility that certain families are more involved in the recovery of lost telomeres than others, which would result in an enrichment near the base of the telomere. On the autosomes, tandemly repeated telomere associated sequences (TAS-L and TAS-R) sit at the base of each telomere. An HTT insertion adjacent to a TAS-L or TAS-R repeat therefore represents the first insertion after the HTT array eroded completely. Similarly, if an HTT has healed a terminal deficiency extending beyond the TAS array, it will form a junction with unique subtelomeric sequence (as described above; see also Kern and Begun (2008)). In both cases, if an HTT family is particularly activated upon the complete erosion of a telomere, it will be enriched next to these subtelomeric sequences relative to the other HTT families. We find that *HeT-A2* and *TART* family elements are underrepresented and enriched, respectively, at the telomere base. The most striking result though is that *TAHRE* is 4-fold underrepresented at the base (Figure 6D). Since *HeT-A* and *TAHRE* likely share the same transposition machinery, one would expect them to show the same pattern. Our results therefore suggest the possibility that *HeT-A* elements are more proficient at using *TAHRE’s* transpositional machinery to heal telomeres than *TAHRE* itself. Regardless, our data suggest that the HTTs differ in their propensity to heal completely eroded telomeres, and some (particularly the *TART* family) may be especially activated by telomere loss.

## DISCUSSION

Linear chromosomes require a specialized mechanism to replicate and maintain their ends, and most eukaryotes use the highly conserved telomerase enzyme to do so. Yet Drosophila and a few other groups have dispensed with telomerase and instead use the HTT retrotransposons to perform this essential function, highlighting the remarkable flexibility of evolution (Pardue and DeBaryshe 2008). Extensive efforts have provided full assemblies of several telomeres in *Drosophila melanogaster* and other Drosophila species (Saint-Léandre *et al*. 2019; George *et al*. 2006). Other studies have hinted that there may be extensive variation in telomere length and structure within populations (Kern and Begun 2008; Wei *et al*. 2017), but in the absence of a large-scale population genomics analysis, it remains difficult to distinguish among very different evolutionary models that could govern telomere dynamics.

While long-read sequencing technologies are beginning to provide glimpses into structural variation of complex repetitive regions (Chang *et al*. 2019; Weissensteiner *et al*. 2017), the vast majority of population genomics data comes from short-read Illumina sequencing. Here, we leveraged the ConTExt pipeline and available population genomic data to investigate HTT variation in *D. melanogaster* at an unprecedented scale, examining 85 lines derived from five world-wide populations and encompassing close to 10,000 HTT insertions.

### Caveats and technical considerations

There are some limitations resulting from the data we used that are derived from pools of ∼50 inbred whole females of varying ages. First, the GDL lines have continued to evolve under inbreeding and laboratory evolution, which could result in increased variance of TE copy number, and fixation or loss of certain TE variants. However, this also would apply to most other studies of Drosophila telomeres in which strains have not been freshly collected from the wild before sequencing, including reference genome assembly (Saint-Léandre *et al*. 2019; George *et al*. 2006). Although not necessarily reflecting natural variation, inbreeding and continued lab evolution may provide increased power to detect TEs escaping repression or differences in their susceptibility to repression. Second, telomere length might differ across tissues and over an individual’s lifespan. However, this is unlikely to affect our estimates as 1) these differences should be specific to individual flies and 2) germline erosion is reported to be approximately 70 bp per generation in Drosophila (Levis 1989; Biessmann *et al*. 1990a), much lower than the variance we report here. Some of our inferences about rates and patterns of telomere variation could in principle be extended using a mutation accumulation framework, although the relatively low per-generation rates would require very large sample sizes. Finally, a powerful, albeit extremely costly, way to further probe variation in telomere structure would be to directly assemble full telomeres from wild populations, as Saint-Léandre *et al*. (2019) have reported from single strains of multiple Drosophila species using long-read sequencing.

The ConTExt method infers the presence and arrangement of TEs from paired-end Illumina data. We validated ConTExt here by examining five genomes for which both Illumina and PacBio assembly data are available (Long *et al*. 2018), and simulating Illumina short reads from the PacBio assembly. We found a high concordance for HTT copy number estimates between the simulated and actual ConTExt analyses (Figure 1C).

We have further confidence in our analyses based on the concordance of our results with a wide range of previous studies. For example, the large fraction of truncated HTTs we observed is consistent with the assembled telomeres from the *D. melanogaster* reference strain (George *et al*. 2006). In addition, the high prevalence of tandem HTTs is similar between the two studies. For sequence diversity, our estimates for TE families across the 85 GDL concord with estimates reported from analyses of the ISO-1 reference strain (Kaminker *et al*. 2002). We also identified a large number of terminal deficiencies, consistent with a more limited previous study (Kern and Begun 2008). When examining copy number, we found a strong correlation between the number of *HeT-A* family and *TAHRE* elements in outlier strains, which supports prior suggestions that *TAHRE* is the autonomous regulator of HTT transposition (Abad *et al*. 2004). Finally, our identification of *R*-elements inserted outside of the rDNA agrees with previous studies (Stage and Eickbush 2009).

### Frameworks for considering HTT evolution

HTTs are often considered to be a canonical example of transposon domestication, perhaps reflecting an adaptive response to ameliorate a reduction in telomerase activity (Mason *et al*. 2008; Pardue and DeBaryshe 2011; Arkhipova 2012; Shpiz and Kalmykov 2012; Servant and Deininger 2015). Others though have noted the rapid evolution of HTTs and telomere-associated proteins and proposed that HTTs remain in genetic conflict with their hosts (Saint-Léandre *et al*. 2019; Lee *et al*. 2017; Cosby *et al*. 2019; Markova *et al*. 2020). Markova et al. (2020) have also proposed recently that telomeric localization of HTTs might reflect a strategy of site-specific integration to minimize deleterious effects on the host, rather than domestication. They also noted that the HTTs are likely to maintain the ability to cause genetic conflict by evolving through new mutations or gene conversion. We suggest that a fourth factor, niche instability, is an additional and under recognized factor that influences HTT evolution. We outline and examine our results below in the context of these four distinct processes that may affect telomere dynamics in Drosophila.

### Evidence for domestication or host selection

The pioneering essays on selfish and junk DNA suggested that repetitive DNAs might occasionally be co-opted by their hosts to serve essential functions (Doolittle and Sapienza 1980; Orgel and Crick 1980). Active TEs will experience continued selection for their own replicative capacity and so complete domestication likely requires ablating transpositional activity (Jangam *et al*. 2017). Consistent with this, most described examples of domesticated TEs have lost the ability to transpose. In contrast, the HTTs have not. The extent of HTT domestication is a challenging question to address with population genomic data, as it likely relates to the ways in which their activity is regulated by the host genome, how this regulation is distinct from that of other TEs, and how dependent they have become upon telomere-specific host factors. The elaborate system of HTT control also speaks to host domestication. Like other TEs, the HTTs are controlled by the piRNA pathway, though unlike most TEs it is the active elements themselves that act as the piRNA clusters (Shpiz and Kalmykova 2012). This potentially makes it challenging for the host to exert stable control of telomeres since the piRNA source is at an inherently unstable location. HTTs outside of the telomere could thus indicate evidence of host control. We suggest that the heterochromatic, likely inactive, *TAHRE* tandem array we discovered in 82/85 strains could be a locus of host control, functioning as either a source of *TAHRE* piRNAs or producing dominant-negative *TAHRE* proteins.

Full domestication of HTTs would imply that the HTT transposition rate does not increase beyond a point where telomere length becomes excessive for host fitness. Although there is clearly extensive variation in telomere length, there are only limited data on whether and how telomere length may affect fitness (Walter *et al*. 2007). While our analysis thus does not provide direct evidence for domestication, the lack of full domestication would cause genetic conflict, which we discuss next.

### Evidence for genetics conflict

Typical TEs exist in conflict with the host genome, as they gain a replicative advantage through transposition but also deleteriously impact host fitness by disrupting functional sequence, facilitating ectopic recombination, and creating intrachromosomal breaks (Kelleher *et al*. 2020). HTTs insert specifically at the ends of chromosomes and so avoid causing these harmful impacts. If long telomeres have no negative fitness impacts, then adoption of this insertional strategy may have largely resolved the typical TE-host conflict. On the other hand, if excessive elongation causes decreased host fitness, conflict of two sorts arises: 1) conflict with the host genome and 2) conflict among the HTTs competing for limited insertion sites.

First, while there should exist an optimal rate of telomere elongation for chromosome integrity, higher rates of transposition may be optimal for the HTTs as they are for TEs in general, with faster replicating subfamilies replacing slower ones (Charlesworth and Langley 1986). This conflict with the host should skew telomeres toward longer-than-optimal lengths and require the evolution of host suppression to shift the rate of elongation closer to the host’s optimum. The optimal average telomere length for *D. melanogaster* is unknown, so this potential skew cannot be detected from population data. However, we observed several strains with exceptionally long telomeres, the longest of which also exhibited extreme CN expansions of several non-telomeric TEs, suggesting heterochromatin maintenance or piRNA pathway defects. The extent of telomere elongation in these strains highlights the degree to which host suppression limits HTT replication, and the extent to which HTTs expand in its absence, suggestive of conflict. As noted above though, these relatively rare strains with significant TE expansions might reflect laboratory evolution of the strains rather than true natural variation.

Second, there is potential conflict among HTTs, as they are all restricted to the same genomic region. This should favor the emergence of variants with higher rates of transposition, displacing less active variants. While we found some differences in the proportion of HTT subfamilies between long and short telomere strains, the three major families are equally represented, suggesting that longer telomeres result from loss of host control rather than runaway expansion of one family at the expense of others. We therefore do not have evidence of strong genetic conflict among HTTs.

Conflict with either the host or other HTTs should lead to the fixation of new sequence variants within the HTTs, and ongoing conflict may result in substantial diversification. It is tempting, therefore, to infer conflict from the number of distinct HTT subfamilies and the sequence diversity within them. However, our observation that the copy number variation of most HTT SNPs is consistent with recently active elements, in contrast to typical TEs where most SNPs are consistent with older insertions, suggests that the rapid turnover of the telomeres has a major influence on the patterns of HTT diversification. This distinctly high rate of sequence loss must be accounted for when trying to infer evolutionary forces influencing HTT evolution (see below).

### Evidence for niche specialization of HTTs and tradeoffs

One route by which TEs might attenuate conflict with the genome is by limiting their deleterious impacts, for example by inserting solely at gene-poor loci. Both the HTTs and R-elements appear to have adopted such strategies, inserting at gene-poor but unstable loci, thereby trading long-term insertion sites for reduced host fitness costs (Markova *et al*. 2020). TE subfamilies could conceivably reacquire a more general insertion pattern to escape both the instability and competition with the other TEs targeting the same limited set of occupiable sites. We observed no clear evidence of HTT insertions outside of the telomere, but identified 33 strain-specific *R1*-element insertions outside of the rDNA. Multispecies surveys of *R*-elements and HTTs have provided similar observations, with low to moderate fractions of *R*-element being inserted outside of the rDNA in most species (Stage and Eickbush 2007) but few, if any, non-telomeric HTT insertions (with the possible exception of *D. rhopaloa*, Saint-Léandre *et al*. 2019). This could reflect domestication locking the HTTs more tightly into their insertion site than *R*-elements, perhaps through reliance on telomere proteins. But a neutral explanation based on TE mobilization/transposition mechanism is also conceivable. *R*-elements have a functional endonuclease which is required to insert in the rDNA. *R*-elements therefore only need to relax the sequence specificity of their endonuclease to escape the rDNA. In contrast, it is not clear if the HTTs depend on endonuclease to insert at the chromosome ends, even if *TART* and *TAHRE* elements retain a potentially functional endonuclease domain (Morrish *et al*. 2007; Arkhipova 2012). If HTTs do not utilize an endonuclease as part of their normal lifecycle, then escape from the telomeres likely requires reacquisition of a functional endonuclease, thus locking telomeric TEs more tightly into their insertion site than the *R*-elements.

We observed that HTTs have a propensity to form tandem insertions of the same subfamily, which might reflect another aspect of niche specialization that mitigates some of the fitness cost of being exposed to terminal erosion. *Het-A* insertions are known to include short tags of sequence derived from the upstream element whose 3’-end-promoter initiated transcription, which George *et al*. (2010) suggest provides a buffer to protect the element from erosion. We note that the tandems we describe are 1) generally between elements of the same *HeT-A* subfamily and 2) the tandems present in the reference genome include arrays of full-length elements (Fig 6C), rather than a single full-length element preceded by short fragments. We suggest, therefore, that these tandems reflect sequential insertion of the same subfamily to the chromosome end. The most terminal HTT on a telomere is at imminent risk of loss by erosion. An HTT that generates two or three tandem copies at the same chromosome end exposes the distal-most insertion to immediate erosion but buffers the more proximal elements for potentially hundreds of generations. By contrast, creating two independent copies on two different telomeres generates the same number of new insertions, but both are subject to immediate erosion. For the *HeT-A* family and *TAHRE*, this is potentially an evolved strategy of niche specialization, because they do not have an inherent mechanism such as internal repeats to catalyze tandem formation. *TART* family elements do, however, so their occurrence in tandem is more likely to reflect a mutational process, as we discuss below.

### Evidence for mutational processes and genome instability driving HTT turnover

The erosion and replacement of sequence is a fundamental characteristic of telomeres, and occupying such an unstable locus thus distinguishes the HTTs from most other TE families. Our observations both clarify the extent of this instability and highlight its impact on HTT evolution. First, the presence of terminal repeats in *TART* family elements predisposes them to tandem expansions by ectopic recombination, similar to LTR retrotransposons. The high rate of tandem *TART* elements with shared terminal repeats we observed is evidence of recombination-mediated changes in telomere length. This extends upon prior observations of *TART* family tandems in the ISO-1 reference genome (George *et al*. 2006), and highlights that not at all gains and loss of sequence result from transposition and erosion. Second, we discovered a high prevalence of terminal deletions in the GDL, extending previous discoveries (Levis 1989; Walter *et al*. 1995; Kern and Begun 2008). Surprisingly, we also discovered that about one third of terminal deletions are unhealed by an HTT. It is possible that unhealed deletions are more common in inbred lab stocks; for example, if inbreeding allows alleles to fix that reduce HTT activity. Regardless though, most telomeres are presumably eventually healed over evolutionary time, and this creates opportunities for new HTT subfamilies to expand.

Third and most generally, we find that for positions within HTT sequences that display multiple alleles, both alleles tend to exhibit high copy number variation consistent with being recently active elements. This contrasts sharply with other active TEs for which only one allele generally appears to be recently active while the other typically shows low copy number variation, reflective of being older and likely inactive. We suggest this results from HTT insertions being lost much more rapidly in the telomeres than other TEs elsewhere in the genome, where the relics of past transposition may be preserved for millions of years. In the telomeres, the window into the past is much more narrow, due to continuous erosion. Therefore, the diversity of HTT families and subfamilies does not necessarily demonstrate rapid evolution for HTTs, but may instead simply reflect that each species is recreating its entire population of HTT insertions all the time as erosion and recombination erase older insertions. This will be reflected in their phylogenetic pattern, as the HTTs show common ancestors with long terminal branches (Saint-Léandre *et al*. 2019; Villasante *et al*. 2007), rather than a birth/death pattern with very short terminal branches such as for active primate LINE-1 elements (Khan *et al*. 2006).

### Conclusion

HTTs are often described as a clear-cut case of domestication. They clearly serve an essential host function, and non-telomeric insertions appear to be negligible. But niche specialization is a plausible alternative, whereby telomere localization reflects an evolutionary strategy of the HTTs to reduce their fitness impact on the genome (Markova *et al*. 2020). In this interpretation, HTTs may have become the dominant mechanism for forming telomeres as a by-product of evolving preferential localization to telomeres. Species such as silkworm might represent a transitional stage, as they have telomere-associated retrotransposons while maintaining telomerase (Okazaki *et al*. 1993, 1995; Pardue and DeBaryshe 2008).

On the other hand, genetic conflict has been suggested to be a major driver of the observed rapid evolution of HTT families and subfamilies, telomeric proteins, and subtelomeric sequences (Saint-Léandre *et al*. 2019; Lee *et al*. 2017; Cosby *et al*. 2019; Saint-Léandre and Levine 2020). But direct evidence for conflict is elusive, largely due to limited data on fitness consequences of telomere length variation. We further suggest that the high instability of chromosome ends is an alternative explanation of HTT variability. However, our results are also consistent with prior observations that host sequences regulate the HTTs and further suggest that their susceptibility to regulation varies among the HTTs and is actively evolving, consistent with ongoing conflict between these genomic endosymbionts and their host genome. We conclude that multiple evolutionary forces and mechanistic processes interact to explain the patterns of HTT variation, with telomere instability being an under-recognized contributor.

## MATERIALS AND METHODS

### The repeat index and nomenclature

We used the manually curated repeat index described in McGurk and Barbash (2018), which contains consensus sequences for the known *D. melanogaster* TE families as well as satellite repeats, including the left and right telomere associated sequences (TAS-L and TAS-R). The telomeric TE sequences in this index are somewhat distinct from the Repbase entries (Jurka *et al*. 2005). First, the long perfect near-terminal repeats (PNTR) found in the *TART* family elements are given entries separate from the internal sequences, similar to how the terminal repeats of LTR retrotransposons are given their own entries separate from the internal sequence in Repbase. Second, the index includes four *HeT-A* family entries in addition to the Repbase consensus (Table S1), as many of the *HeT-A* family insertions in the reference genome are quite divergent from the consensus sequence (< 85% identity, Figure S4A). To ensure reads derived from *HeT-A* family insertions would align to the repeat index, we extracted all insertions annotated as *HeT-A* in the UCSC genome browser Repeat Masker track for the release 6 *D. melanogaster* genome. We then performed all pairwise alignments using BLASTn and constructed a graph by construing insertions as nodes and connected all pairs of sequence that shared >90% identity with edges (McGurk and Barbash 2018). We partitioned this graph into communities of homologous insertions using the Louvain algorithm (Blondel *et al*. 2008). This identified four *HeT-A* family communities, one of which appeared to contain subcommunities. We split the large community into two subcommunities by reclustering with the alignment cutoff set at 95% identity. From each community we manually chose a full-length element as representative and added these to the repeat index, labelling them *HeT-A1* to *HeT-A5*. However, we removed *HeT-A4* due to high sequence similarity with *Het-A3*. We refer to the original *HeT-A* sequence from RepBase as “*HeT-A*” (Bao *et al*. 2015). The final set of *Het-A* family sequences all have less than 85% identity to each other, with most pairs having roughly 80% (Figure S3A, Supplemental File 1). For TEs that have internal repeats, we removed them by scanning along each consensus, and for each 70-mer in the consensus masking all subsequent 70-mers within 5 mismatches.

### Categorizing TE families as active

We categorize TE families as active or inactive based on their percent identity and population frequency as summarized in Kelleher and Barbash (2013). We consider a family as putatively active (or recently so) if the mean pairwise identity between individual insertions in the reference genome is greater than 95% (Kaminker *et al*. 2002; Kelleher and Barbash 2013) and the average population frequency of insertions is less than 0.4 (Kofler *et al*. 2012). We additionally consider the telomeric TEs and the two *R*-elements as being active, as well as including *P*-element, *Hobo*, and */*-element. While this approach is likely to misclassify some truly active elements as inactive, the two categories should be fairly representative of the active and inactive families in *D. melanogaster*.

### Sequence data and read mapping

Sequencing data come from a previously published study in which pools of ∼50 females from 85 wild-derived *Drosophila melanogaster* lines covering five continents were sequenced with Illumina 100 nt paired-end reads to an average depth of 12.5X (Grenier *et al*. 2015, NCBI BioProject PRJNA268111).

We employed the ConTExt pipeline to organize these data and discover structural and sequence variation within the repetitive portions of these genomes (McGurk and Barbash 2018). Briefly, ConTExt aligns repeat-derived reads to the corresponding repeat consensus sequences (using Bowtie2 v. 2.1.0, Langmead and Salzberg 2012). Mixture modeling is used to infer the set of underlying structures that generated the set of discordantly aligned read pairs in each sample, and ConTExt subsequently clusters these junctions to determine which structures are present in multiple samples. We modified the pipeline here to infer sequence variation within repeat families from the aligned reads (described below). Additionally, rather than align reads first to individual insertions and then collapse these alignments onto consensus sequences (McGurk and Barbash 2018), here we aligned reads directly to the consensus sequences using permissive alignment parameters (`--score-min L,0,-2.5 -L 11 -N 1 -i S,1,.5 -D 100 -R 5`).

### Filtering ambiguous alignments

While Bowtie2’s mapping quality summarizes the ambiguity of alignments, it heavily penalizes divergence from the reference sequences and is defined in a rather opaque manner involving nested conditionals. Because we expect some reads to derive from TE insertions diverged from the consensus, it is undesirable to penalize reads with no secondary alignments but which are nonetheless diverged from the consensus. We instead filter ambiguous alignments by directly considering the primary and secondary alignment scores, which we convert to percent identities assuming all penalties are due to mismatches. We use these to define a score *M* reflecting the distance between the primary (*AS*) and secondary (*XS*) hits:

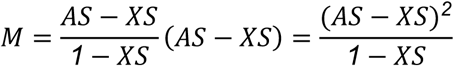

This score summarizes the distance between the primary and secondary alignments and is penalized by the divergence of best alignment from the consensus, but it does so in a predictable fashion. If a secondary alignment is reported by Bowtie2, we require this score to be greater than 0.05 for the alignment to be considered unambiguous and included in the analysis. If the primary hit perfectly matches the consensus, the secondary alignment must be more than 5% diverged from the consensus for the read to be included in the analysis. If the primary alignment is 10% divergent, the secondary alignment must be more than 20% diverged from the consensus for the read to be included in the analysis (Figure S3B). Finally, we exclude any read whose primary alignment is more than 20% diverged from the consensus.

### Overview of statistical analyses

We describe the details of each model employed in the subsequent sections, but outline the general logic of our analyses here. We approached the analyses of the data from the perspective of Bayesian parameter estimation, seeking to define probability distributions over all possible values of our statistics of interest to guide our interpretations. To accomplish model-checking, we employ posterior predictive simulations to evaluate the ability of our models to account for features of the observed data as a means of model checking. When the failure of a model to account for features of the data is an observation of interest, we report the posterior predictive p-value (Gelman 2013), which unlike the frequentist p-value is conditioned upon the set of models most consistent with the data rather than a predefined null-hypothesis.

When relating read counts to copy number, we employ the negative binomial distribution as it can model overdispersed counts and the sum of its random variables are also negative binomially distributed, allowing us to describe the read count distributions under different copy numbers. That is, if the read depth of a single copy structure is negatively binomially distributed with mean *μ* and dispersion *α*, we can model the read depth *x* of a structure present in *n* copies as *x* ∼ *NB*(*nμ, nα*). We further employ partial pooling, which jointly estimates the copy number of individual structures and the underlying copy number distribution from which junctions arise, using the complete set of observed read counts to constrain the individual copy number estimates and prevent outlier read counts from being unduly interpreted as multicopy structures. When making inferences about copy number distributions, we focus on identifying the mean and variance parameters of the distributions. However, TE copy number, and consequently telomere length, are prone to outliers relative to a normal distribution, even in log scale, which likely reflect atypical events that resulted in a copy number expansion. Therefore, we model the copy number distribution as a mixture of two distributions, one reflecting typical copy number variation and the other reflecting the distribution after extreme copy number expansion, and incorporate latent variables that assign each observed copy number as an inlier or an outlier.

When estimating relative quantities such as proportions or frequencies we employ beta-binomial and Dirichlet-multinomial models, and whenever we believe that strain-specific factors, such as telomere composition, are relevant, we incorporate this hierarchical structure into the model.

For Bayesian modelling we employ PyMC3 (v. 3.9). For each model, we draw samples from the posterior distribution in two chains to assess convergence, sampling continuous random variables with the No-U-Turn Sampler initialized with jitter+adapt_diag as implemented in PyMC3. The No-U-Turn Sampler (NUTS) is the current state of the art in Monte Carlo sampling, capable of efficiently exploring high-dimensional parameter spaces and drawing uncorrelated samples from the posterior. While superior to the Metropolis algorithm in most regards, it is unable to sample discrete random variables due to its reliance on gradient information. Consequently, we represent copy number in our analyses with positive continuous random variables rather than with discrete variables, preferring greater confidence in the posterior sample afforded by NUTS. We note that treating copy number as continuous is not dissimilar from the commonly used approach of estimating copy number by dividing the observed read counts by the expected read depth. Further, as the genomic DNA for each GDL strain was obtained from a pool of individuals, the average copy number per individual in the stock is better described by a continuous rather than discrete variable. Similarly, when we incorporate outlier detection in our models we marginalize out the binary labels representing the outlier status of individual observations to avoid the need for Gibbs sampling. We retain the interpretability provided by these labels by using the posterior distribution to reconstruct the probability that each observation reflects an outlier.

### Modelling read depth

To infer the copy number of multicopy sequence, we need to know the read depth distribution of single-copy sequences. We estimate this from coverage of the two major autosomes in the reference genome, using the same filtering steps described in McGurk and Barbash (2018). As in McGurk and Barbash (2018), we consider the coverage of sequence not by the reads themselves, but rather by the interval between read pairs. However, here we model the read depth of single-copy sequence with negative binomial distributions, allowing the mean and overdispersion to vary with %GC. To better handle junctions with extreme %GC, we model the relationship between the mean and overdispersion of %GC in each strain *j* with library-specific functions:

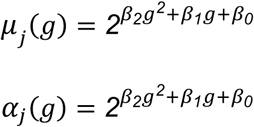

where *g* is the expected %GC of read pairs spanning a position and the *β*s are the coefficients of each quadratic function (Figure S3D, D).

Additionally, there is a clear excess of positions with zero coverage in the data that is likely due to filtering, but homozygous deletions may also contribute. We account for this by fitting a zero-inflated negative binomial distribution when inferring the mean and overdispersion functions, to ensure the excess of zeroes does not downwardly bias the expected read depth. We allow the amount of zero-inflation to vary by %GC, relating it with a logistic function as the expected proportion of zeros must be restricted between 0 and 1:

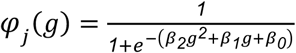

For each sample we obtain maximum-likelihood estimates of these functions using PyMC3.

### Estimating copy number from read depth

For each position of a consensus sequence we count the number of concordant read pairs which span it in each strain. We estimate the number of copies containing a given position by dividing the read count at each position by the expected read depth given the model of GC-bias we inferred for that strain (Figure S3C, D). From this we estimate sequence abundance by summing the copy number of each position. The edges of the sequence have reduced mappability as most read pairs span a junction between the repeat and some other sequence. To account for this, we exclude the first and last 500 bp from our estimate of sequence abundance and divide the resulting sum by 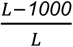, where *L* is the length of the consensus sequence. We do not estimate sequence abundance for consensus sequences shorter than 1,000 bp.

When estimating the total amount of HTT sequence comprising the telomeres, we do not filter reads for ambiguous alignments, as most ambiguity in reads aligning to the HTTs is caused by homology among related subfamilies, so while it may be unclear which HTT gave rise to such a read it is almost certainly derived from the telomere. However, when estimating the copy number of specific HTT families or subfamilies, we do filter out ambiguously aligned reads.

### Estimating the breakpoints of terminal deficiencies from read depth

We identified terminal deficiencies by visually examining the coverage of subtelomeric sequence for obvious loss of read depth (Supplemental File S3). To estimate the breakpoint of a terminal deficiency based on read depth, we sought to identify a sequence coordinate beyond which the copy number, estimated as the ratio between the observed and expected coverage, drops below 1. For each of the autosome arms, we considered coverage of the most telomere-proximal 100 kb sequence and thinned it by retaining only every hundredth position, to account for autocorrelation in the coverage signal. We then removed any position that had more than five repeat-masked nucleotides within 100 bp to reduce the impact of repeat masking on coverage. At each position, we divided the observed read count by the expected read count given the local GC-content. We then fit a step function to these copy-number estimates, *x*, where the expected copy number differs from 1 on the telomere-proximal side of a breakpoint, *c*. We estimate this breakpoint by minimizing the sum of squared residuals (SSR), defined for positions on the centromere proximal side of the breakpoint as

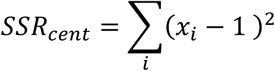

and on the telomere-proximal side as

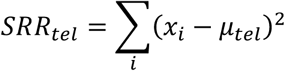

where *μ*_*tel*_ is the mean copy number estimate on the telomere-proximal side of the breakpoint. We combine these as

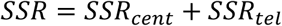

We noted a tendency for the last 10 kb of the 2R subtelomere to have reduced coverage in most strains, which may result from decreased mappability. We were therefore conservative when calling deficiencies in this region, requiring a complete loss of read depth to call a deficiency in the absence of an HTT-subtelomere junction, and only calling heterozygous deficiencies if a HTT-subtelomere junction accompanied the drop in coverage.

If we had evidence of an HTT-subtelomere junction at a breakpoint, we used the sequence coordinates of the junction as the deficiency breakpoint. In estimating the number of independently derived deficiencies, we considered any pair of deficiencies within 200 nt of each other as potentially the same allele and collapsed them.

### Analysis of truncated HTTs

HTTs, like all non-LTR retrotransposons, are frequently 5’-truncated due to incomplete reverse transcription, in addition to being susceptible to telomere erosion. While ConTExt does not permit the reconstruction of HTT insertions, we can determine the extent to which each insertion is 5’-truncated. A full-length insertion must have the intact 5’-end, though the presence of the 5’-end does not guarantee that there are not deletions within the insertion. We note, however, that we observe few junctions consistent with internal deletions within HTTs. Given the high rate of truncation shown in Figure 2C, strains containing only truncated insertions of a particular family might be common. But because such elements are unlikely to encode for functional proteins, and only the autonomous *TAHRE* and *TART* family elements encode reverse transcriptase, some full-length elements are likely maintained by selection. We found that 33% of strains lack any full-length *TART* family elements, but only 2 out of 85 strains lack full-length *TAHRE*. To assess the likelihood of this observation, we model the number of full-length elements and then used posterior predictive simulations to ask whether we observe fewer strains without any full-length *TAHRE* elements than would be expected under the fitted model.

We estimate the length of insertions by considering the coordinates of an element’s 5’-end in an HTT-HTT junction, and define an element as being full-length if its estimated end is within 300 bp of the expected 5’-end of its consensus sequence. As elements of the *TART* family are flanked by terminal repeats, we require the junction to involve the 5’-end of the PNTR rather than the internal sequence to be considered full-length, though some fraction of such junctions may involve the 3’-PNTR of *TART* instead and reflect truncated elements. As the counts are overdispersed, for each family we model the number of full-length insertions, *k*_*i*_ out of all *N*_*i*_ insertions in strain *i* with a beta-binomial distribution. For interpretability, we estimate mean, *q*, and concentration, *v*, parameters that respectively describe the expected binomial rate of full-length elements and the variability around this which drives the overdispersion. We define the priors in this model as

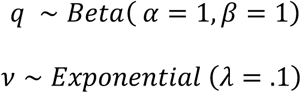

placing a uniform prior on the rate of full-length elements. We model the likelihood as

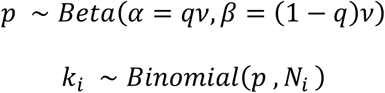

To assess the extent to which the model accounts for the fraction of strains lacking any full-length elements, we employ posterior predictive simulations to obtain a posterior p-value that describes the probability of observing as few or fewer strains without full-length elements present in the GDL given the posterior distribution. In 99% of the simulations, the fitted model predicted more strains entirely missing full-length *TAHRE* than we observed. One interpretation is that selection acts on the genome to maintain the reverse transcriptase encoded by *TAHRE*. Alternatively, in most genomes there may be a full-length *TAHRE* insertion in tandem with another HTT that is rarely deleted, perhaps because it is located outside of the telomere. We present in the Results evidence in favor of this alternative explanation, but stress that these *TAHRE* elements are old and likely non-functional.

### Interpreting sequence variation within repeats

In our analyses of sequence variation, we estimate the copy numbers of the different alleles found at a given position. We first count the number of reads supporting each allele at that position, excluding nucleotides whose PHRED base quality score is less than 30 as well as those in the first and last five bases of the read. This filtering score reduces the contribution of sequencing and alignment errors, but decreases the read count. Therefore, allele copy number estimates based on these counts are downwardly biased. To infer the copy number of these alleles, we therefore also compute the proportion of reads that support each allele and multiply by the estimated copy number at that position. If the copy number of an allele is estimated to be a small fraction (< 0.2), we assume it reflects sequencing errors and treat it as zero.

To estimate the sequence diversity at a position, we pool these allele copy numbers across all strains in the GDL, treating all copies of the family as members of the same population of TEs. We then estimate diversity as

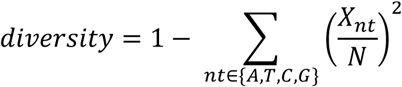

where *N* is the total copy number at that position and *X*_*nt*_ is the copy number of a given allele. We define the major allele as the allele with the greatest copy number in the GDL and the minor allele as that with the second greatest copy number. We estimate the mean and variance of these alleles from their estimated copy number in each strain.

### Correcting for additional biases in read counts over junctions

In our analyses of the read counts over HTT-HTT junctions we often observe fewer reads than expected given a single-copy junction, which could reflect residual heterozygosity, the presence of additional sequence within the junctions such as 3’-tags or polyA-tails, or reads having been filtered out due to alignment ambiguity. We could determine the zygosity status of the junctions between HTTs and healed terminal deficiencies independently of their read counts by examining the coverage of unique sequence on the telomere-proximal side of the junction. Despite that most such deficiencies are truly homozygous, we noted the same deflation of read counts we observed for HTT-HTT junctions, leading us to conclude this reflects a downward bias rather than heterozygosity. To assess the degree of this bias, we modelled the read counts over these HTT-deficiency junctions as

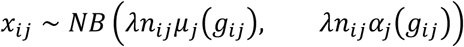

where *n*_*ij*_ is the true copy number of the junctions (1 if homozygous and 0.5 if heterozygous), *μ*_*j*_ (*g*_*ij*_) is the expected read count given the %GC (Figure S3C, D) and *α*_*j*_ (*g*_*ij*_) is the degree of overdispersion. The degree of bias is *λ* over which we place a prior of

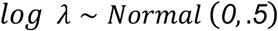

Fitting this model, we find that the read count over these junctions is 43% (95% credible interval 35% - 49%) of what we expected given the read depth distributions observed for single copy sequence. We incorporate this bias (and our uncertainty about its exact degree) into our copy number estimates of junctions between HTTs and other repeats, calibrating their read counts against this subset of HTT junctions with known copy number. When we use these corrected copy number estimates to infer the total amount of telomeric sequence in each strain, we find a strong concordance with our estimates based on consensus coverage without MapQ filtering (Figure 1D).

### Identifying TE copies from their junctions

We identify and count the numbers of euchromatic TE insertions as described in McGurk and Barbash (2018), by identifying the 3’ and/or 5’ junctions the TE forms with its insertion site. For *R*-elements, we estimate the copy number from the read count of the junction between their 3’-end and the rDNA. As all such *R*-element junctions all have the same sequence coordinates and are present in multiple copies rather than being a set of single and low-copy junctions like the HTT-HTT junctions, we estimate their copy as the ratio between the observed and expected read count (assuming a single copy structure with the same %GC) without needing to employ partial pooling to avoid overestimating their copy number. As described in the next section, we employ partial pooling to estimate the copy number of HTT insertions, as we expect there might be both single and multi-copy HTT-HTT junctions.

However, as we estimated the copy number of TE insertions (by counting junctions) in unique sequence differently from that of the HTTs and the *R*-elements (by also considering read counts), we worried that some differences among these categories could reflect differences in the behavior of these estimators rather than true biological differences. To rule this out, we also estimated copy number from the read depth near the 3’-ends of each family’s consensus, estimating copy number as the average sequence abundance over a 1 kb interval ending 500 bp from the element’s 3’-end. Considering only the 3’-end helps limit the impact of 5’-truncation on estimated copy number. While this ensures the copy numbers of all TE families are estimated in the same way, it does not distinguish TE-derived satellites, such as the large *R1*- and *Bari1*-derived satellite arrays, from dispersed TEs. We are, however, able to distinguish these in the copy number estimates based on junctions rather than read depth (i.e. tandem *Bari1* and *R1*-elements form junctions with themselves, rather than with unique sequence or the *R*-element’s rDNA insertion site).

### Modelling copy number of HTT-HTT junctions

While many of the junctions we identify may reflect single-copy structures, it is likely that an appreciable fraction of junctions are present in multiple copies. We therefore estimated the copy numbers of HTT junctions based on their read counts. For a junction *i* in sample *j*, we model its read depth *x*_*ij*_ as arising from a negative binomial distribution truncated at 2, as we only analyzed junctions supported by at least two reads

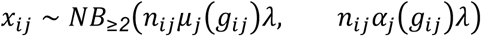

Here *μ*_*j*_ and *α*_*j*_ are the library-specific functions that describe how the read depth and overdispersion vary with the expected %GC of read pairs spanning the junction, and *g*_*ij*_ and *λ* are the aforementioned bias. Finally, *n*_*ij*_ is the quantity of interest: the underlying copy number of the junction.

If we estimated this copy number for each junction independently with weakly informative priors, we would upwardly bias our copy-number estimates. This is because *a priori* we should expect that most HTTs junctions are truly single-copy and that higher than expected read counts often reflect single-copy structures that generated more reads than expected by chance. But, as we do not know ahead of time what fraction are single-copy, we sought to infer the underlying copy number distribution at the same time we estimate the copy numbers of the individual junctions. This accomplishes the parameter estimation analog of a multiple-test correction, by shrinking copy-number estimates away from extreme values unless the data justify believing the copy number greater than 1. We assume that the mode of this copy-number distribution is 1, and represent how much junction copy number varies around this mode with a variance parameters (Figure S2 A, B)

Thus we model the overall copy-number distribution of HTT junctions as

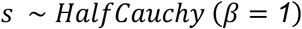

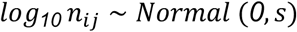

where *s* is the standard deviation of the copy number distribution. We model the reduced read counts over HTT junctions as

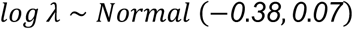

We bring these together to model the read counts over individual HTT junctions with a negative binomial distribution, truncated at 2 to reflect the minimum junction read count for inclusion in this analysis:

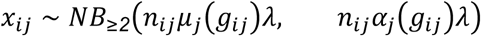

### Estimating the proportion of missed junctions

The GDL strains were sequenced at relatively low depth, ranging from 10-20 reads expected per junction. This, coupled with the filtering out of HTT reads that could not be assigned unambiguously, means that some fraction of junctions truly present in a strain must have been missed by our analysis. We therefore sought to quantify the extent of this. To this end, for each of our 2,000 posterior samples of the read count model’s parameters we simulated new read counts from a non-truncated negative binomial distribution

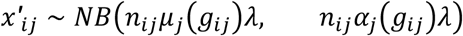

This provided a large set of read counts generated from junctions whose copy number, GC-content, and read depth are well-matched to those observed in the GDL. For each strain in each of the 2,000 replicates of this simulation, we computed that fraction of junctions with fewer than two reads. For each strain, we take the average of this value across all 2,000 as an estimate of the rate at which truly present junctions would be missed.

We note that there may be some downward bias to this estimate, as the set of observed junctions will be more enriched for multicopy junctions than the set of missed junctions. So these simulations assume a higher proportion of multicopy junctions than may have been truly present in the GDL.

### Modelling telomere length and copy number distributions

Telomere length and HTT copy number distributions share similar features, so we model both under the same framework. First, as both are restricted to positive values, we model their log-transformed values. Second, we observe values that are outliers relative to this log-normal distribution. These outliers, for reasons discussed in the Results, almost certainly reflect events that occurred while the strains were maintained as lab stocks. Consequently, we model the data as a mixture of two distributions, one largely reflecting natural variation and the other reflecting anomalous copy number expansions.

To infer the telomere-length distribution of each population in a way that accounts for the observed outliers, we modeled the log_10_-transformed total telomere length *x*_*i*_ of a typical individual *i* in population *j* as arising from a Normal distribution, with population-specific means, *μ*_*i*_, and standard deviation, *σ* _*i*_.

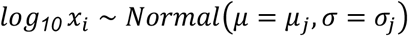

If the observation reflects an atypical copy number expansion we model it as arising instead from

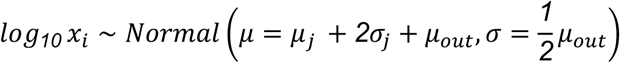

where *μ*_*out*_ reflects how much greater than two standard deviations above the inlier mean the outliers tend to be and over which we place a prior which considers an order of magnitude copy number increase a plausible outlier

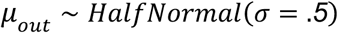

The outlier status of each individual could be modeled with a variable *z*_*i*_which equals 0 for inliers and 1 for outliers so that

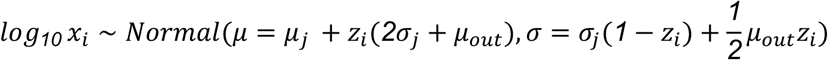

and we would model *z*_*i*_ as being Bernoulli distributed with a probability of being an outlier *p* arising from a uniform distribution on the interval 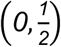

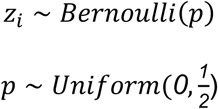

However, as binary variables are challenging to sample we marginalize these labels out of the model (Hogg *et al*. 2010),

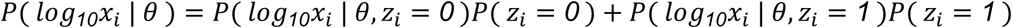

where *θ* represents all model parameters (*μ*_*j*_, *μ*_*out*_, *σ* _*j*_) other than the outlier labels, and *P*(*z*_*i*_ = *1*) = *p* and *P*(*z*_*i*_ = *0*) = *1* − *p*. To compute the posterior probability that an observation is an outlier we use

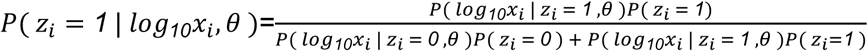

and average over all values of *θ* sampled from the posterior. Those strains that have a probability of being outliers are depicted in Figure 2A and Figure S1B-J as black dots.

When modelling telomere length we describe our prior beliefs as

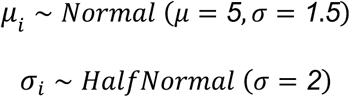

These priors imply that average total telomere lengths may be as small as 1 kb or as large 100 Mb. Similarly, the prior on *σ* is open to the standard deviation of telomere length encompassing four orders of magnitude. Both priors are somewhat overly permissive allowing the data to drive the parameter estimates with little constraint from our prior beliefs.

### Modeling the propensity of different HTTs to heal telomere erosion

To determine whether some families have a higher propensity to heal telomere erosion, we compared the frequency with which an element is found next to subtelomeric sequence against the proportion of HTT copies it comprises in each strain. As junctions with subtelomeric sequence involve the 3’-ends of the HTTs, we estimate the copy number of each family from the number of junctions involving the element’s 3’-end, ensuring any biases resulting from mappability or structural variation at the 3’-end of the elements affect both estimates. We employ a Dirichlet-Multinomial model where the number of junctions *x*_*ij*_ between an HTT family *i* and the known chromosome 2 and 3 telomere associated sequence (TAS) repeats across all strains *j* arises as

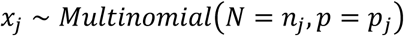

where *n*_*j*_ is the total number HTT-TAS and HTT-deficiency junctions we detect in strain *j, p*_*j*_ is a vector describing the probability of finding each HTT family at the base of the telomere in strain *j*,

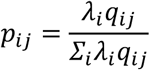

*λ*_*i*_ is the relative enrichment of family *i* at the base of the telomere, and *q*_*ij*_ is the number of HTT elements belonging to family *i* in the strain *j*, incorporating the differences in telomere composition among strains into the model. We model our prior beliefs about the relative enrichment as being wholly agnostic using a uniform Dirichlet prior

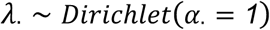

We fit this model considering the HTT families as well as subfamilies.

### Modelling interspersion among HTTs

To assess the tendency of particular HTTs to neighbor each other we also employ a Dirichlet-Multinomial model. Here we consider an interspersion matrix *X* where each cell *X*_*ijk*_ counts the number of times the 3’-end of element *i* forms a junction with 5’-sequence of element *j* in strain *k*. We model the vector of counts in strain *k* as arising from

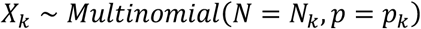

where *p*_*k*_ is a matrix describing the probability that two elements *i* and *j* neighbor each other. This relates to both a tendency, *λ*_*ij*_, of two elements *i* and *j* to neighbor each other and the proportion of HTTs elements *i* and *j* found in strain *k*

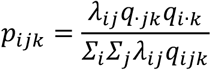

where *q*_·*jk*_ and *q*_*i*·*k*_ are the marginal counts

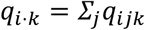

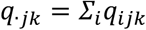

We model prior beliefs about the parameters of interest, the vectorized matrix of associations, with a uniform Dirichlet distribution

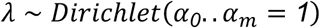

We note that this does not model the interspersion under reorderings of existing telomeres but rather under the generation of new telomeres drawn from the observed proportions.

### Assessing performance on simulated data

We simulated Illumina Hiseq 2000 paired-end data from five available GDL PacBio genome assemblies (B59, I23, N25, T29A, and ZH26; Long *et al*. 2018) using ART (version: MountRanier). To match the characteristics of the GDL NGS data, we set the genome-wide read depth and fragment size distributions to match those reported for the particular GDL strains. We ran these simulated datasets through the ConTExt pipeline using the same parameters as for the real GDL data.

To assess our ability to correctly estimate the copy number of HTT insertions, we compared the copy-number estimates from the simulated data to the copy number evident in the assemblies from which the data were simulated. To determine the ground truth for HTT copy numbers in each assembly, we aligned the HTT consensus sequences to the assemblies using BLAST and filtered out hits with less than 90% identity to a consensus. We then counted the number of hits corresponding to the intact 3’-end of each HTT family in the assembly.

## Data availability

Sequencing data come from a previously published study (NCBI BioProject PRJNA268111, Grenier *et al*. 2015). ConTExt is located at https://github.com/LaptopBiologist/ConTExt. Release 6 of the reference genome is described in Hoskins *et al*. (2015); the 5 PacBio assemblies of GDL strains are described in Long *et al*. (2018).

## Author’s contributions

Conceptualization: M.P.M, A.M.D.C. and D.A.B., Method development: M.P.M., Analyses: M.P.M and A.M.D.C., Interpretation: M.P.M, A.M.D.C. and D.A.B. All authors wrote, read and approved the final manuscript.

## Acknowledgements

We are grateful to Alla Kalmykova, Mia Levine, Homa Papoli and Jesper Boman who commented on an earlier version of this manuscript. We also thank Andrew Clark, Cédric Feschotte, Alexander Suh and the members of their labs for stimulating discussions.

## SUPPLEMENTARY TABLES

**Table S1: HTT copy numbers in Pacbio assemblies and corresponding ConTExt estimates**. This table presents for each HTT family, in each of the five Pacbio assemblies, the copy number estimated from BLAST alignments of consensus sequences against the assembly and the copy number that was estimated from applying ConTExt to Illumina data simulated from each assembly.

**Table S2: TE copy number per strain**. The estimated copy number of each TE family in the 85 GDL strains.

## SUPPLEMENTARY FIGURES

**Figure S1.**
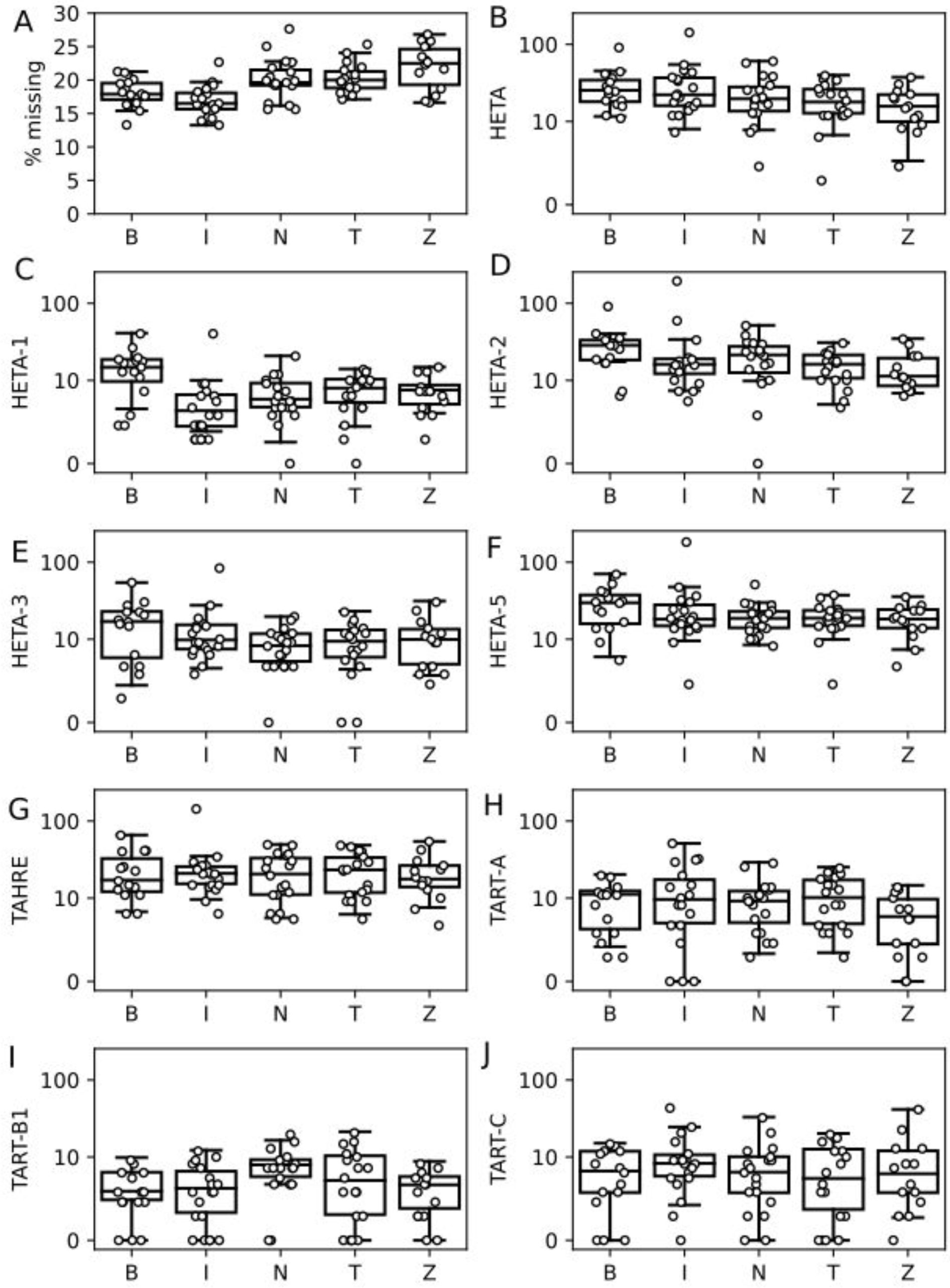
The copy number of the HTT subfamilies in each strain, broken down by population and corrected by the estimated fraction of missing junctions. A) The percent of junctions for all HTT subfamilies inferred from posterior predictive simulations to have been missed in each strain due to read depth variation, broken down by population. To ensure interpretations of population structure are not driven by these missing junctions for the rest of this figure only, when assessing the presence of population structure we roughly corrected our copy number estimates by dividing by one minus the inferred fraction of missing junctions for each HTT subfamily. Inferred missing junctions for each HTT subfamily: B) *HeT-A*, C) *HeT-A1*, D) *HeT-A2*, E) *HeT-A3*, F) *HeT-A5*, G) *TAHRE*, H) *TART-A*, I) *TART-B*, J) *TART-C*.

**Figure S2.**
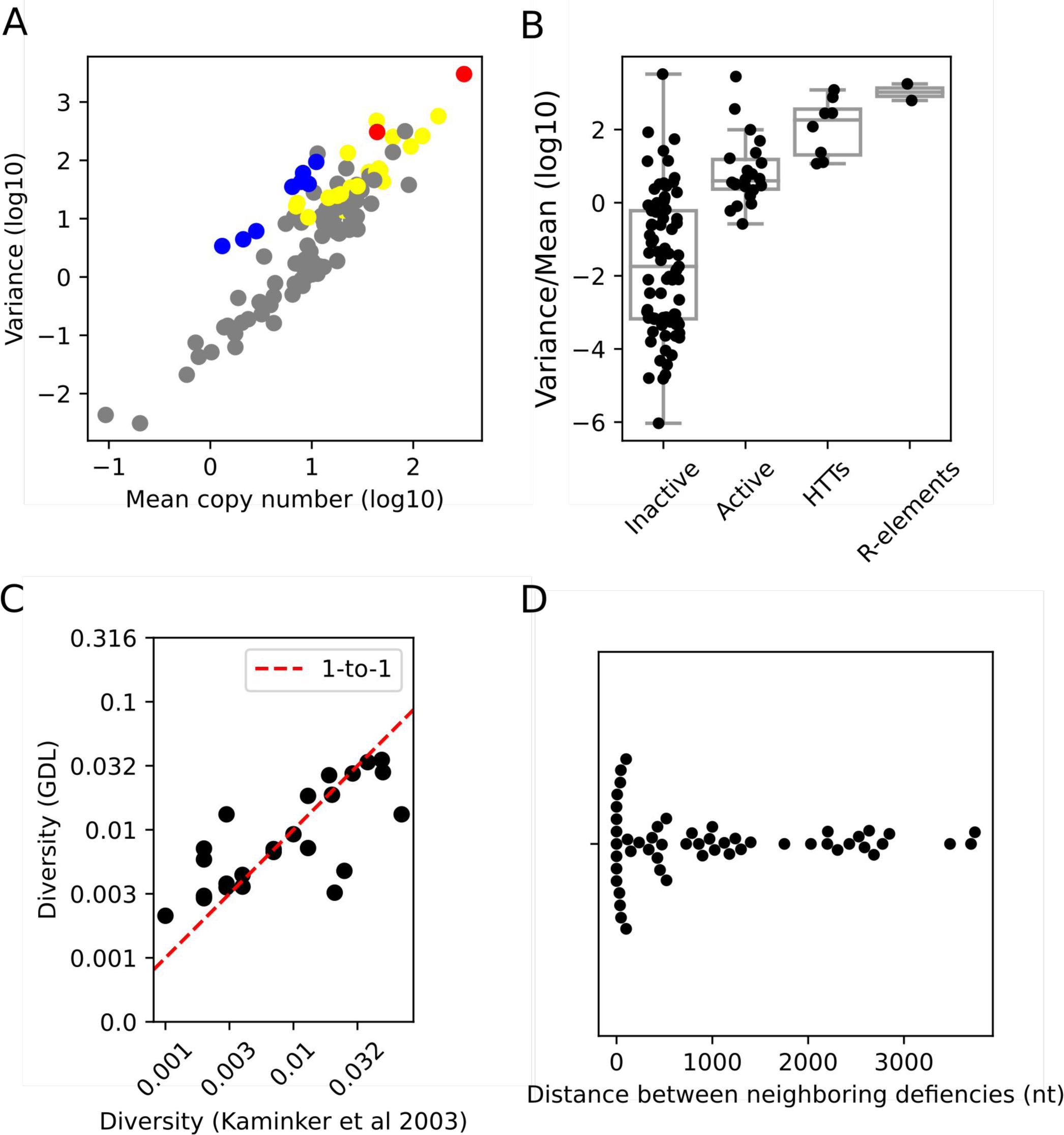
Alternative and supporting visualizations of data. (A) Scatter plot showing the relationship between mean copy number (log scale) and variance (log scale) of TE families across the GDL as estimated from the read depth at the 3’-end of TE consensus sequences rather than junctions. Designations of active and inactive TEs are from prior estimates of sequence divergence and population frequency as described in the Materials and Methods. Solid line represents the expected relationship under the assumption of little variation in population frequency and low linkage disequilibrium among insertions. (B) Boxplots depicting distributions of variance-to-mean ratios (log_10_) for each of the four categories of the TE families based on these copy number estimates. (C) A scatter plot comparing our estimates of sequence diversity for the TE families in Figure 4B with those reported in Kaminker et al (2003). This is restricted to TE families included in both analyses. (D) A swarm plot depicting the distribution of distances between terminal deficiency breakpoints, where for each breakpoint the distance is plotted relative to the nearest other breakpoint we discovered in a different sample.

**Figure S3.**
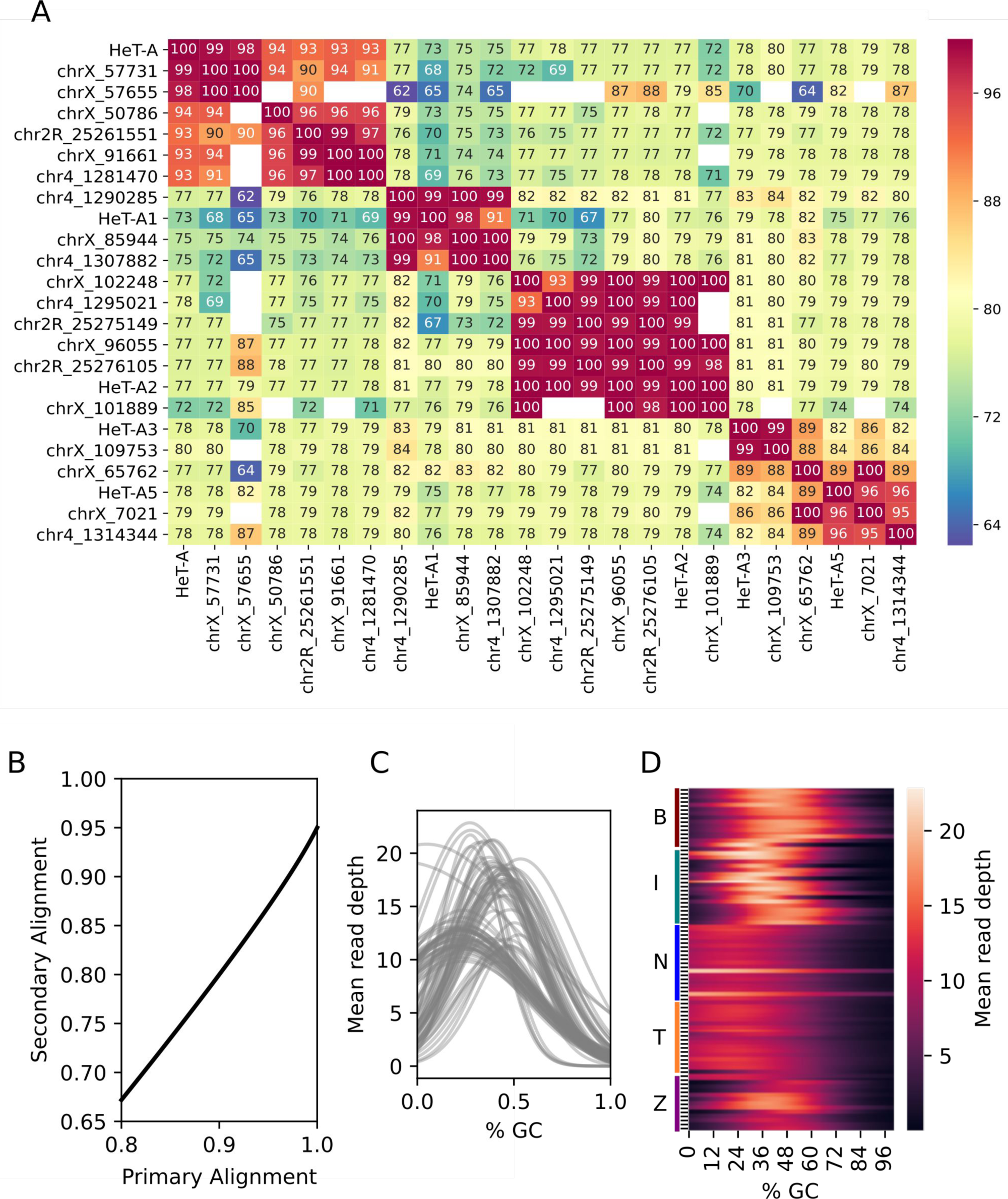
Details on interpreting alignments to telomeric transposable elements. (A) Percent identities of all >300 bp insertions annotated as *HeT-A* in the UCSC genome browser repeat-masker track of the Release 6 reference-genome telomeres. Percent identities are based on a ClustalOmega multiple sequence alignment with default parameters. The Repbase consensus sequence and the representatives we chose for each *HeT-A* subfamily in our repeat index are indicated as such. The white cells indicate no alignment between a pair of sequences, all involving alignments with the two shortest *HeT-A* fragments. (B) A depiction of how we define ambiguous alignments. The black line indicates how diverged the second best alignment must be for a primary alignment with a given percent identity to be considered unambiguous. (C) The expected relationship between read depth and %GC for each of the GDL strains. The variability of this relationship across strains is what necessitated controlling for %GC when estimating copy number. (D) A heatmap of the expected relationship between read depth and %GC, sorted by population.

## SUPPLEMENTARY FILES

**Supplementary File 1. FASTA file of HTT consensus sequences for five *HeT-A* subfamilies (*HeT-A, HeT-A1, HeT-A-2, HeT-A3, HeT-A5*), *TAHRE*, and three *TART* subfamilies (*TART-A, TART-B1, TART-C*)**. The positions of the insertions in the Release 6 *Drosophila melanogaster* reference genome used as representative sequences for the *HeT-A* subfamilies are indicated.

**Supplementary File 2. Plots of mean-variance relationships between allele copy number for each active TE family**. As in Figures 4D-E, each dot reflects the observed mean and variance of the copy number of the major (blue) and minor (gold) alleles of positions with >0.1 sequence diversity. For reference, the shaded regions are re-plotted from Figure 3B.

**Supplementary File 3. Visualization of read coverage on chromosomes with evidence for terminal deficiencies**. Each blue dot is the estimated copy number at a given position, determined as the ratio between the observed and expected read depth. The black lines below the plots indicate which regions of the reference are repeat-masked (and should show little or no read coverage). The orange trendlines are a 10 kb rolling window average of estimated copy number and the red lines indicate the average copy number estimate to the left and right of the inferred deficiency breakpoint. The vertical red line in some plots indicates the location of the HTT-subtelomere junction. Below each plot we note the breakpoint coordinate we estimated based on read depth as well as the percent of the coverage prior to the breakpoint remaining in the deleted regions, from which we infer the heterozygosity or homozygosity of the deficiency.

